# Microplastic consumption induces inflammatory signatures in the colon and prolongs a viral arthritis

**DOI:** 10.1101/2021.08.24.456180

**Authors:** Daniel J. Rawle, Troy Dumenil, Bing Tang, Cameron Bishop, Kexin Yan, Thuy T. Le, Andreas Suhrbier

## Abstract

Global microplastic (MP) contamination and the effects on the environment are well described. However, the potential for MP consumption to affect human health remains controversial. Mice consuming ≈80 µg/kg/day of 1 µm polystyrene MPs via their drinking water for a month showed no weight loss, nor were MPs detected in organs. The microbiome was also unchanged. MP consumption did lead to small transcriptional changes in the colon suggesting plasma membrane perturbations and mild inflammation. Mice were challenged with the arthritogenic chikungunya virus, with MP consumption leading to a significantly prolonged arthritic foot swelling that was associated with elevated Th1, NK cell and neutrophil signatures. Immunohistochemistry also showed a significant increase in the ratio of neutrophils to monocyte/macrophages. The picture that emerges is reminiscent of enteropathic arthritis, whereby perturbations in the colon are thought to activate innate lymphoid cells that can *inter alia* migrate to joint tissues to promote inflammation.

## INTRODUCTION

Global plastic production has increased exponentially and was ≈259 million metric tons in 2018 (Hirt and Body-Malapel, 2020). When polluted into the environment, these plastics fragment and degrade into microplastics (MPs) (WHO, 2019)(Dawson et al., 2018). MPs are now extremely widespread in our environment (Lau et al., 2020) spanning multiple biomes from the ocean floor (Amaral-Zettler et al., 2020) to near the top of Mt Everest (Napper et al., 2020), with our epoch being referred to as the age of plastics, the Plasticene (Campanale et al., 2020). We now even have the plastisphere, with a unique biotope associated with plastic pollution (Amaral-Zettler et al., 2020). The extensive presence of MPs in our environment means the opportunities for consumption by humans is widespread (Cox et al., 2019), with MPs readily detectable in human colons and feces (Ibrahim et al., 2021; Schwabl et al., 2019). Some beverage items have high levels of MPs, for instance, plastic tea bags (Hernandez et al., 2019), disposable plastic-lined paper cups (Ranjan et al., 2021), some bottled water (Jin et al., 2021b; Wong et al., 2021), and infant formula prepared in polypropylene feeding bottles (Li et al., 2020b). Many foods are contaminated with MPs (Jin et al., 2021b) including bivalves (Garrido Gamarro et al., 2020), table salt (Zhang et al., 2020) and instant rice products (Dessi et al., 2021). MPs are also found in the air and are breathed in (Akanyange et al., 2021) and after mucociliary clearance would be swallowed (Vethaak and Legler, 2021). Even the simple act of opening plastic containers is associated with MP exposure (Sobhani et al., 2020).

The question of whether and how MPs might affect human health remains controversial, with considerable speculation and a shortage of compelling data (Lim, 2021; Vethaak and Legler, 2021). Although difficult to determine, human MP consumption has been estimated to be ≈175 µg - 9 mg/kg/day (de Wit and Bigaud, 2019; Senathirajah et al., 2021). Studies on the effects of MPs *in vivo*, primarily in mice have not produced compelling data (Böhmert et al., 2019; Braeuning, 2019; Deng and Zhang, 2019), with a range of often contradictory results reported (Hirt and Body-Malapel, 2020; van Raamsdonk et al., 2020). Mice are generally fed polystyrene (occasionally polyethylene) MP beads ranging in size from 0.5-1 µm (Deng et al., 2018; Luo et al., 2019a; Stock et al., 2019) to 150 µm (Li et al., 2020a). The approximate MP doses (converted to weight/kg/day), range from ≈15 µg/kg/day (Jin et al., 2019; Lu et al., 2018; Luo et al., 2019a; Luo et al., 2019b) to ≈20 mg/kg/day (Deng et al., 2017; Li et al., 2020a; Stock et al., 2019; Yang et al., 2019), and reach as high as 100 mg/kg/day (Deng et al., 2020). In some studies MPs are found in visceral organs (Deng et al., 2017; Jin et al., 2019), whereas others find no uptake into the body, even at high doses (Stock et al., 2019). Some studies reported weight loss in mice after MP consumption (Jin et al., 2021a; Lu et al., 2018; Xie et al., 2020), whereas others did not (Deng et al., 2017; Deng et al., 2018; Hou et al., 2021; Luo et al., 2019a; Luo et al., 2019b; Stock et al., 2019; Wang et al., 2021b). Reported adverse effects of MP consumption include disturbance of lipid/energy metabolism (Deng et al., 2017; Deng et al., 2018; Jin et al., 2019; Lu et al., 2018; Luo et al., 2019a; Luo et al., 2019b; Zheng et al., 2021), oxidative stress (An et al., 2021; Deng et al., 2017; Xie et al., 2020), microbiota dysbiosis (Jin et al., 2019; Li et al., 2020a; Lu et al., 2018), reproductive toxicity (Jin et al., 2021a; Luo et al., 2019a; Luo et al., 2019b; Park et al., 2020; Xie et al., 2020), overt intestinal perturbations (Jin et al., 2019; Li et al., 2020a; Luo et al., 2019b; Zheng et al., 2021) and inflammation (Li et al., 2020a; Wang et al., 2021b; Zheng et al., 2021). In contrast, others found no inflammation, oxidative stress or other adverse health effects (Stock et al., 2019). Although difficult to unravel, some key sources of artefactual results might include (i) avoidance by mice of MP-containing water because it tastes or smells tainted (Janssens et al., 1995; Villberg et al., 2004; Villberg et al., 1997) leading to dehydration and weight loss, (ii) trauma caused by repeated oral gavage (Kinder et al., 2014), (iii) cage effects leading to artificial microbiome heterogeneity (Basson et al., 2020), (iv) unrealistically high MP doses that might cause fecal impaction (Issac and Kandasubramanian, 2021) or other gut damage (Yong et al., 2020) and/or (v) the presence of sodium azide in most commercial sources of plastic beads, with this compound known to be toxic to mammals and cause oxidative stress (Tat et al., 2021).

We and others have previously described in detail a wild-type mouse model of infection and disease caused by the arthritogenic alphavirus, chikungunya virus (CHIKV) (Suhrbier, 2019). Mouse infections result in a reproducible robust transient viraemia, joint infection and an overt self-resolving arthritic foot swelling associated with a prodigious, predominantly mononuclear inflammatory infiltrate (Gardner et al., 2010; Prow et al., 2019; Prow et al., 2017; Wilson et al., 2017). CHIKV arthritic inflammation is predominantly driven by Th1 CD4 T cells and to a lesser extent NK cells (Nakaya et al., 2012; Suhrbier, 2019; Teo et al., 2015), with increased neutrophils associated with promotion of arthritis in various settings (Poo et al., 2014a; Prow et al., 2019). Monocytes/macrophages have a role both in promoting viral arthritides and in resolution of inflammation (Felipe et al., 2020; Labadie et al., 2010; Poo et al., 2014a; Prow et al., 2019). We chose to challenge mice that had consumed MPs with CHIKV because this infection results in widespread transcriptional changes (Wilson et al., 2017), providing potential insights into the effects of MPs on *inter alia*the prodigious anti-viral responses and the multifactorial, self-resolving, arthritic immunopathology (Suhrbier, 2019).

Herein we have addressed the aforementioned potential sources of artifacts in mouse MP studies, and use RNA-Seq to show that MP consumption leads to a mild proinflammatory signature in colon, which was associated with exacerbation of a viral arthritic immunopathology. Overall the study supports the notion that MP consumption can exacerbate immunopathology

## Results

### 1 µM MPs in drinking water do not enter mouse tissues

In order to provide a detailed insight into the effects of MP consumption in mammals *in vivo*, C57BL/6J mice were given MPs in their drinking water. The MPs compromised 1 µM Sephadex beads internally loaded with a fluorescent dye, with 10^6^ MP/ml (0.526 µg/ml) added to the drinking water. The beads were purchased sodium azide-free and were washed prior to use.

MP-supplementation did not affect water consumption, water preference, food consumption or weight gain (Supplementary Fig. 1a-d). Over a month, mice consumed on average 3.85 ± SD 0.31 ml water per day per mouse, and 3.84 ± SD 0.095 ml of water plus MP per day per mouse (Supplementary Fig. 1a). MP consumption was thus estimated to be 3.84 x10^6^ MP/day/mouse or 2.02 µg/day/mouse, which converts to a MP dose of ≈80 µg/kg/day (Fig. 1a). The human consumption of MP has been estimated to be 0.1 - 5 g per week (de Wit and Bigaud, 2019; Senathirajah et al., 2021), which converts to a dose of ≈175 µg - 9 mg/kg/day. The oral mouse MP dose used herein is thus comparable with the lower end of the estimated MP consumption in humans, although clearly 1 µM Sephadex beads do not recapitulate the large size range or diversity in chemical composition of MPs consumed by humans (Banerjee and Shelver, 2021; Campanale et al., 2020). Our dose of ≈80 µg/kg/day is similar to doses used in some previous rodent studies (Jin et al., 2019; Lu et al., 2018; Luo et al., 2019a; Luo et al., 2019b), but is considerably lower than, for instance, the >20 mg/kg/day doses used in other studies (Deng et al., 2020; Deng et al., 2017; Li et al., 2020a; Stock et al., 2019; Yang et al., 2019).

**Figure 1.**
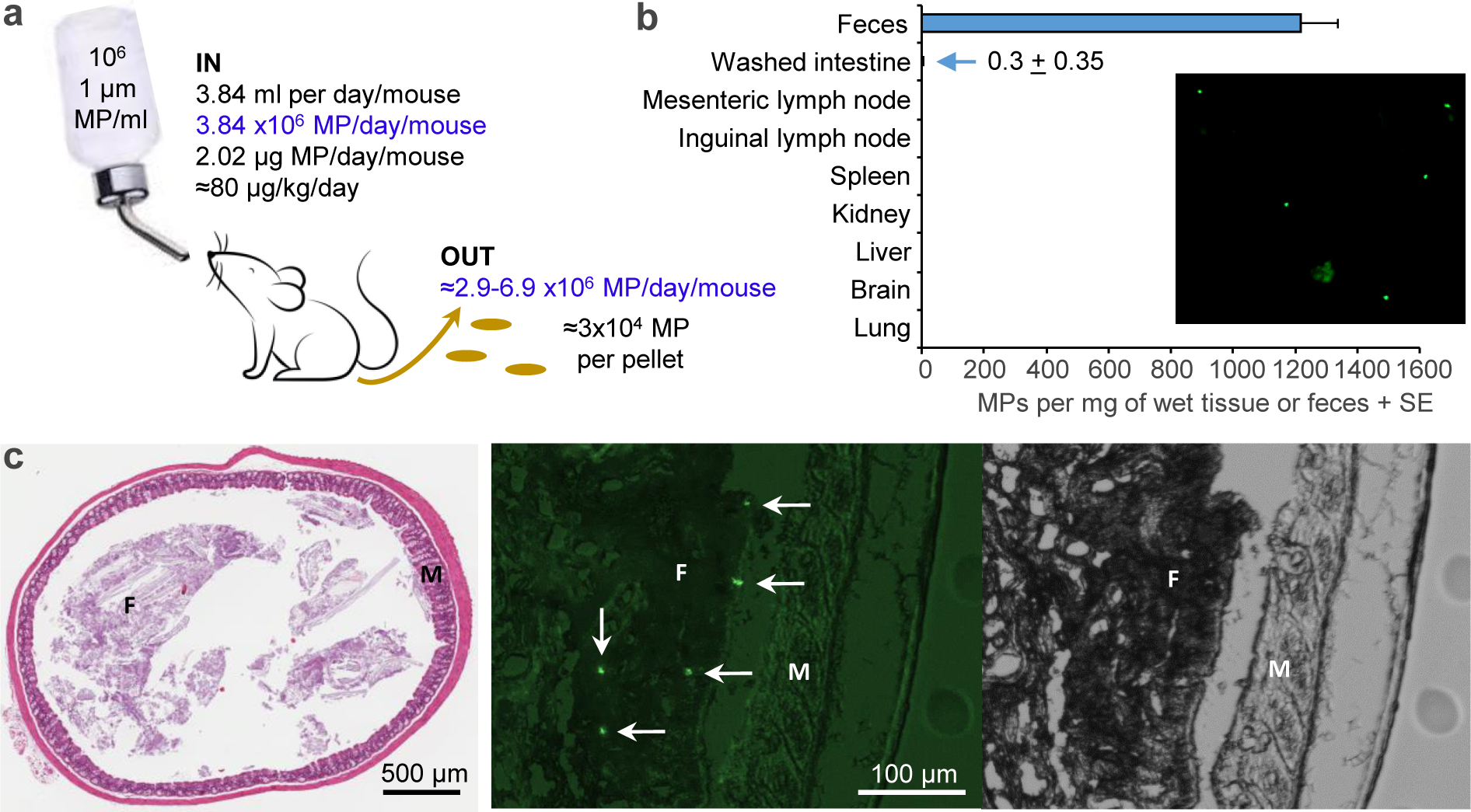
Detection of MP *in vivo*. **a** Mice were given drinking water supplemented with 10^6^ MP/ml for 8 weeks and MP consumption measured by weighing of water bottles. Calculations for MP excretion in fecal pellets derived from data in b. **b** After 8 weeks of MP consumption the indicated organs and fecal pellets were collected (n=3-6 mice per sample), weighed (wet weight) and digested with SDS and proteinase K. MPs in the digests were counted using a haemocytometer and a fluorescent microscope; the inserted florescent image shows MPs from a digest of a fecal pellet. **c** Formalin fixed colon after 8 weeks of MP consumption. H&E staining showing orientation of sectioning across the colon (left). An image from a frozen section (7 µm) of colon viewed under a fluorescent microscope (middle), with a phase contrast image of the same region (right). MPs are indicated by arrows; M - mucosa, F - fecal material. More fluorescent images are shown in Supplementary Fig. 2.

Whether ingested MPs can, under physiological conditions, traverse the gut lining and enter the body to any significant degree remains a matter of some speculation (Braeuning, 2019; Campanale et al., 2020) and may be dose dependent (Stock et al., 2019). Digests of fecal pellets from mice drinking MP-supplemented water indicated that there were ≈1200 MP/mg of feces (Fig. 1b). As mice produce ≈100-240 fecal pellets per day (de Lucia and Ostanello, 2020; Hoibian et al., 2018), this coverts to ≈2.9-6.7 x10 MP excreted per day and ≈3 x10 MP per pellet (mean 24 mg/pellet) (Fig. 1a). This calculation argues that of the ≈3.8x10 MP consumed per day, most if not all are excreted, perhaps arguing against the concept of bioaccumulation (Mohamed Nor et al., 2021). Consistent with this observation, we were unable to detect any 1 µM fluorescent MPs in digests of major internal organs and draining lymph nodes after 8 weeks of drinking MP-supplemented water (Fig. 1b). Some MP were found in digests of washed intestines (Fig. 1b), although whether this was due to <100% effective washing or MP having entered intestinal tissues cannot be determined using this assay. A previous mouse study administering ≈500 µg/kg/day of 1 µM MPs by oral gavage, similarly found no MP in major internal organs (Stock et al., 2019).

Frozen sections of colon from mice that had been drinking MP-supplemented drinking water for 8 weeks were viewed by florescent microscopy. Although MP could readily be seen in feces, no MP were seen in intestinal tissues (Supplementary Fig. 2). Most common bacteria are approximately 1-2 µM in diameter and also do not ordinarily cross the intestinal epithelium (Fine et al., 2020). Taken together with data in Fig. 1b, these results argue that any adverse effects associated with MP consumption identified herein are unlikely due to MPs within intestinal tissues or other organs.

### RNA-Seq analysis shows transcriptional changes in the colon after MP consumption

Previous studies in mice have reported gene expression and histological changes in colons of mice fed MPs, although MP doses were often considerably higher than those used herein (Jin et al., 2019; Li et al., 2020a; Lu et al., 2018). To determine whether more physiologically relevant doses of MPs effect the colon *in vivo,* mice were provided with (+MP) and without MPs (-MP) in their drinking water (Fig. 1a) for 33 days. Mice were then euthanized and colons removed and analyzed by RNA-Seq (Supplemental Fig. 1a). The full gene list is provided in Supplementary Table 1a, with a PCA plot showing a clear segregation of +MP and –MP groups (Supplementary Fig. 3). Differentially expressed genes (DEGs) are provided in Supplementary Table 1b (n=281, q<0.05), with a heat map shown in Fig. 2a.

**Figure 2.**
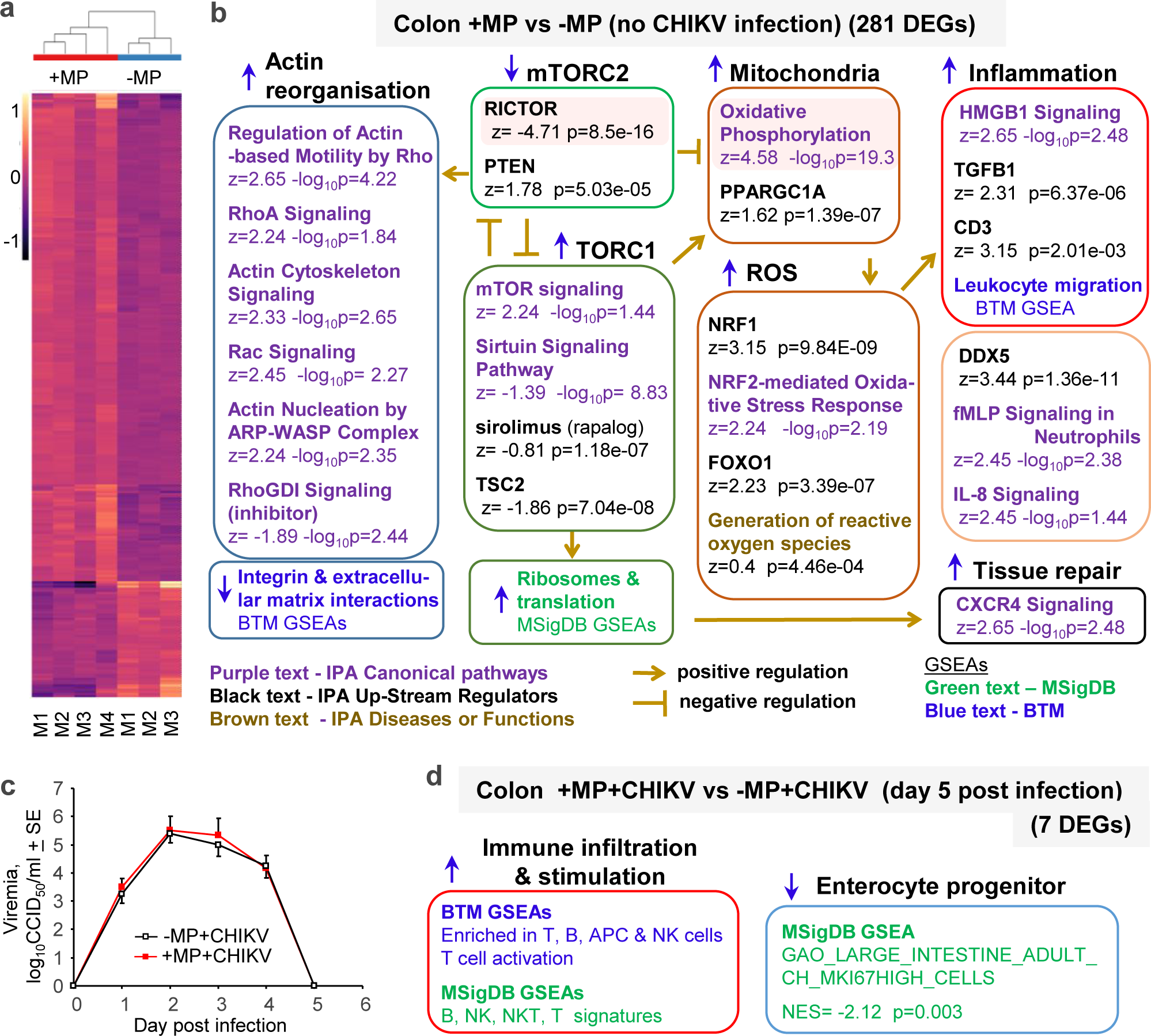
RNA-Seq of colon from mice drinking water +MP versus –MP before and after CHIKV infection. **a** RNA-Seq was used to compare colons from mice drinking MP (+MP) with colons from mice drinking water without MP (-MP) for 28 days. DEGs (n=281) were identified (Supplementary Table 1b) and expression levels for each gene and for each mouse shown as a heat map of the ratio of normalised (DESeq2) log2 expression levels relative to the mean of log2 normalised expression levels for each gene across all mice. **b** The DEGs (n=281) were analyzed by Ingenuity Pathway Analysis (IPA) (direct and indirect); black text – IPA Up-Stream Regulators (USRs) (full list in Supplementary Table 1c), purple text - IPA Canonical pathways (full list in Supplementary Table 1d) and brown text - IPA Diseases or Functions annotation. Light pink shading indicates the dominant highly significant pathways. The full +MP versus - MP gene list (Supplementary Table 1a) was also interrogated by Gene Set Enrichment Analyses (GSEAs) using genes sets from; green text - Molecular Signatures Database (MSigDB) (full list in Supplementary Table 1e) and blue text - Blood Transcription Modules (BTMs) (full list in Supplementary Table 1g). Upward pointing blue arrows indicate up-regulation and downward pointing blue arrows indicate down-regulation. Brown arrows indicate positive regulation, and brown Ts indicate negative regulation. **c** Mice were infected with CHIKV and colons were harvested on day 5 post infection for RNA-Seq analysis. The mean viremias are shown for the mice used in the +MP+CHIKV vs –MP+CHIKV RNA-Seq comparison for day 5 colon. **d** The full gene list for +MP+CHIKV vs –MP+CHIKV (Supplementary Table 2a) was interrogated using GSEAs and BTM and MSigDB gene sets (Supplementary Table 2c,d). Top BTM and selected MSigDB signatures are shown (Supplementary Table 2c,d, highlighted in yellow).

The MP consumption described herein thus caused significant transcriptional changes in the colon, although overall fold changes were low (log_2_ fold change ranged from 1.14 to -1.99). The exact mechanism whereby MPs cause such changes remains unclear, although interaction between the MPs and the lumen side of the colonic epithelium may be involved as (i) MPs were not found in the colonic tissues or in the body (Fig. 1b-c), (ii) MPs can effect human colorectal adenocarcinoma cell line (Caco-2) cultures, with these cells providing an *in vitro* model of the intestinal epithelium (Banerjee and Shelver, 2021; Huang et al., 2021; Stock et al., 2019) and (iii) previous mouse studies have suggested MP consumption can affect the mucosal epithelium and its barrier function (Jin et al., 2019).

### MP consumption results in a mild inflammatory signature in the colon

Ingenuity Pathway Analyses (IPA) of the aforementioned 281 DEGs identified RICTOR as highly significantly down-regulated Up-Stream Regulator (USR) in the +MP group (Fig. 2b, light pink shading; Supplementary Table 1c). RICTOR is a key component of the mammalian Target Of Rapamycin Complex 2 (mTORC2), with active mTORC2 associated with the plasma membrane and mitochondria (Fu and Hall, 2020). The factors that regulate mTORC2 are not well understood (Fu and Hall, 2020; Riggi et al., 2020). However, in yeast hyperosmotic shock, compressive membrane stress and loss of plasma membrane tension can inhibit TORC2 (Le Roux et al., 2019; Morigasaki et al., 2019), suggesting MP interaction with the plasma membrane of the mucosal epithelium is involved. PTEN may play a role (Fig. 2b, mTORC2) as PTEN generates PI(4,5)P2 from PI(3,4,5)P2 (Naderali et al., 2018) and local accumulation of PI(4,5)P2 inhibits TORC2 (Riggi et al., 2020). PI(4,5)P2 also binds N-WASP providing a focus for actin polymerization (Katan and Cockcroft, 2020; Li et al., 2020c) (for actin signatures see below). mTORC1 and mTORC2 counter-regulate each other (Fu and Hall, 2020), with a number of pathway annotations suggesting mTORC1 activity is increased (Fig. 2b, TORC1). Pathway annotations for the drug, sirolimus (an analogue of rapamycin), and for tuberous sclerosis complex 2 (TCS2) have negative z-scores and both specifically inhibit mTORC1 (Huang and Manning, 2008). The Sirtuin Signaling Pathway was significantly down-regulated (Fig. 2b, TORC1), with sirtuin 1 (SIRT1) also an inhibitor of TORC1 (Chen et al., 2018; Ghosh et al., 2010). Gene Set Enrichment Analyses (GSEAs) using gene sets from the Molecular Signatures Database (MSigDB) (Subramanian et al., 2005) also indicated up-regulation of ribosome and translational activity in the +MP group (Supplementary Table 1e), with mTORC1 a key positive regulator of ribosome biogenesis and activity (de la Cruz Lopez et al., 2019; Petibon et al., 2021) (Fig. 2b, green text).

The most significant Canonical pathway identified by IPA was up-regulation of Oxidative Phosphorylation (Fig. 2b, Mitochondria, light pink shading; Supplementary Table 1d).

PPARGC1A, the master regulator of mitochondrial biogenesis, was also up-regulated (Fig. 2b, Mitochondria; Supplementary Table 1c). Up-regulation of mitochondrial activity was supported by Cytoscape analysis (Supplementary Table 1f), as well as GSEAs using gene sets from MSigDB (Supplementary Table 1g) and Blood Transcription Modules (BTMs) (Li et al., 2014) (Supplementary Table 1h). Increased mitochondrial activity has been associated with down- regulation of mTORC2/TORC2 activity in a wide range of settings (Colombi et al., 2011; Heimbucher et al., 2020; Oh et al., 2017; Riggi et al., 2020; Watson et al., 2019), with mTORC1 a well-known positive regulator of mitochondrial biogenesis and functions (de la Cruz Lopez et al., 2019). Increased mitochondrial activity increases production of reactive oxygen species (ROS), with several signatures indicating that MP consumption increased the levels of ROS (Fig. 2b, ROS). NRF1 and NRF2 sense oxidative stress and activate anti-oxidant defenses (Schultz et al., 2014; Zhu et al., 2019). In epithelial tissues ROS also activate FOXO1, leading to induction of anti-oxidant and repair activities (Graves and Milovanova, 2019).

Inflammation signatures were evident in colons of mice consuming MPs (Fig. 2b, Inflammation). Some of these can be associated with elevated levels of ROS (Fig. 2b, Inflammation, top box). HMGB1 requires oxidation for secretion and is a central pro-inflammatory mediator that is also involved in tissue healing and regeneration (Chen et al., 2016; Kwak et al., 2020; Li et al., 2013; Son et al., 2020). TGFB1 is abundantly expressed in the intestine and is up-regulated in inflammatory bowel disease (Ihara et al., 2017), and its activity is positively regulated by ROS (Liu and Desai, 2015; Zhang et al., 2019). A CD3 signature suggested activation of T cells and or NKT cells (Brailey et al., 2020), with ROS well known to stimulate a range of proinflammatory activities including T cell activation (Franchina et al., 2018; Murphy and Siegel, 2013; Yarosz and Chang, 2018). Other inflammation signatures (Fig. 2b, Inflammation, bottom box) included DDX5, which was recently shown to promote intestinal inflammation (Abbasi et al., 2020). IL-8 is a key neutrophil attractant in IBD (Kessel et al., 2021; Singer and Sansonetti, 2004) and N-Formylmethionine-leucyl-phenylalanine (fMLP) mimics the neutrophil chemoattractant and activating activity of a bacterially-derived family of formylated peptides that are believed to promote inflammatory bowel diseases (Somasundaram et al., 2013; Tsai et al., 2016). CXCR4 (the receptor for CXCL12) is involved in tissue repair, with the heterocomplex CXCL12:HMGB1 more active than CXCL12 alone (Bianchi and Mezzapelle, 2020).

A number of IPA pathway annotations indicated that MP consumption promoted actin reorganization (Fig. 2, Actin reorganization). mTORC2 has been shown to be a key regulator of actin organization and remodeling in a wide range of settings including perturbations in the plasma membrane, with this activity involving Rho GTPases such as RhoA and Rac (Diz-Munoz et al., 2016; Li et al., 2020c; Sato et al., 2016; Wu et al., 2016; Xie et al., 2018; Yasuda et al., 2015). GSEAs using BTM gene sets also indicated down-regulation of genes associated with integrin interactions and the extracellular matrix (Fig. 2, Blue text), with integrins and the actin network closely linked in a series of cellular activities (Romero et al., 2020).

### CHIKV infection, weight, viremia and RNA-Seq of colon

Mice drinking water supplemented with (+MP) or without MP (-MP) for 28 days were infected with CHIKV. Mice from both +MP+CHIKV and –MP+CHIKV groups lost a small amount of body weight (≈3%) by day 2 post infection, with mice in the +MP+CHIKV group recovering their weight significantly faster than mice in the -MP+CHIKV group (Supplementary Fig. 4a). The +MP+CHIKV and –MP+CHIKV groups showed similar levels of viremia; this was true both for the mice from which colons were harvested for RNA-Seq analyses (Fig. 2c) and (ii) for two additional independent repeat experiments (Supplementary Fig. 4b).

RNA-Seq analysis for day 5 colons (+MP+CHIKV vs –MP+CHIKV) (Supplementary Fig. 4c Supplementary Table 2a) provided only 7 DEGs (Supplementary Table 2b). GSEAs using BTM gene sets identified multiple dominant signatures associated with T, B, antigen presenting cells (APCs) and NK cells (Fig. 2d; Supplementary Table 2c). GSEAs using MSigDB gene sets contained related signatures, but also identified (as the signature with the lowest NES score) a gene set associated with proliferating (Ki67^hi^) enterocyte progenitors (Gao et al., 2018) (Fig. 2d, Supplementary Table 2d). Five days after CHIKV infection, MP consumption thus appears to have promoted a mild increase in lymphocyte infiltrates in the colon. The mild reduction in enterocyte renewal also suggests MP consumption was somehow protective, reducing the need for tissue repair.

### Low viral loads in the colon and a CHIKV-induced goblet cell response

The CHIKV levels in the colon on day 5 post infection (as measured by qRT PCR and viral titrations) were quite variable, with many samples showing no detectable virus (Supplementary Fig. 4c,d). This variability may have contributed to the inability to identify many DEGs in colon for the RNA-Seq analysis of +MP+CHIKV vs –MP+CHIKV. Viral loads were perhaps marginally lower for the +MP+CHIKV group; however, this was not significant (Supplementary Fig. 4c,d).

A goblet cell signature was identified with a negative z-score (using MSigDB gene sets) for day 5 colon +MP+CHIKV vs –MP+CHIKV (Supplementary Table 2d), with reduced colon goblet cell staining previously reported after MP consumption (Lu et al., 2018). However, AB/PAS staining and Aperio pixel count image analysis, showed no significant effect of MP consumption on goblet cell staining (Supplementary Fig. 5). In contrast, CHIKV infection (i.e. - MP+CHIKV vs +MP, and +MP+CHIKV vs -MP) did show a significant change in AB/PAS staining indicating hyperplasia of goblet cells (Supplementary Fig. 5), which can often be seen after enteric infections (Kim and Khan, 2013).

### Prodigious CHIKV infection signatures in the colon despite low infection levels

RNA-Seq analyses of both +MP+CHIKV vs +MP and -MP+CHIKV vs +MP, showed a series of prodigious CHIKV-induced responses (Supplementary Table 3). These responses may overwhelm (by magnitude and variability) any mild effects associated with MP consumption, perhaps again contributing to the low number of DEGs for +MP+CHIKV vs -MP+CHIKV (n=7).

The CHIKV responses in the colon showed highly significant correlations with responses previous reported for CHIKV-infected feet (Wilson et al., 2017) (Supplementary Fig. 6). When day 5 colon RNA-Seq reads were mapped to the CHIKV genome, 0 to 0.0005% of reads mapped to the viral genome. This is represents a relatively low level of infection when compared with feet, where up to 8% of reads mapped to the CHIKV genome (Wilson et al., 2017). The high level of correlation (Supplementary Fig. 6) despite the low viral loads in the colon, may (at least in part) be explained by responses arising from the circulating proinflammatory cytokines induced by this systemic infection (Gardner et al., 2010).

### Mild inflammation and leukocytosis signatures in MLN after MP consumption

RNA-Seq analysis of mesenteric lymph nodes (MLN) for +MP vs –MP provided only one DEG (Supplementary Fig. 7a; Supplementary Table 4a). MLNs from mice taken 33 and 41 days after initiation of MP consumption were combined in a single analysis (Supplementary Fig. 7a).

GSEAs using BTM and MSigDB gene sets suggested the +MP group had signatures associated with increased inflammation and increased leukocytes (Fig. 3a; Supplementary Table 4b, c); findings consistent with the increased inflammation and leukocyte migration signatures identified in colon (Fig. 2b).

**Figure 3.**
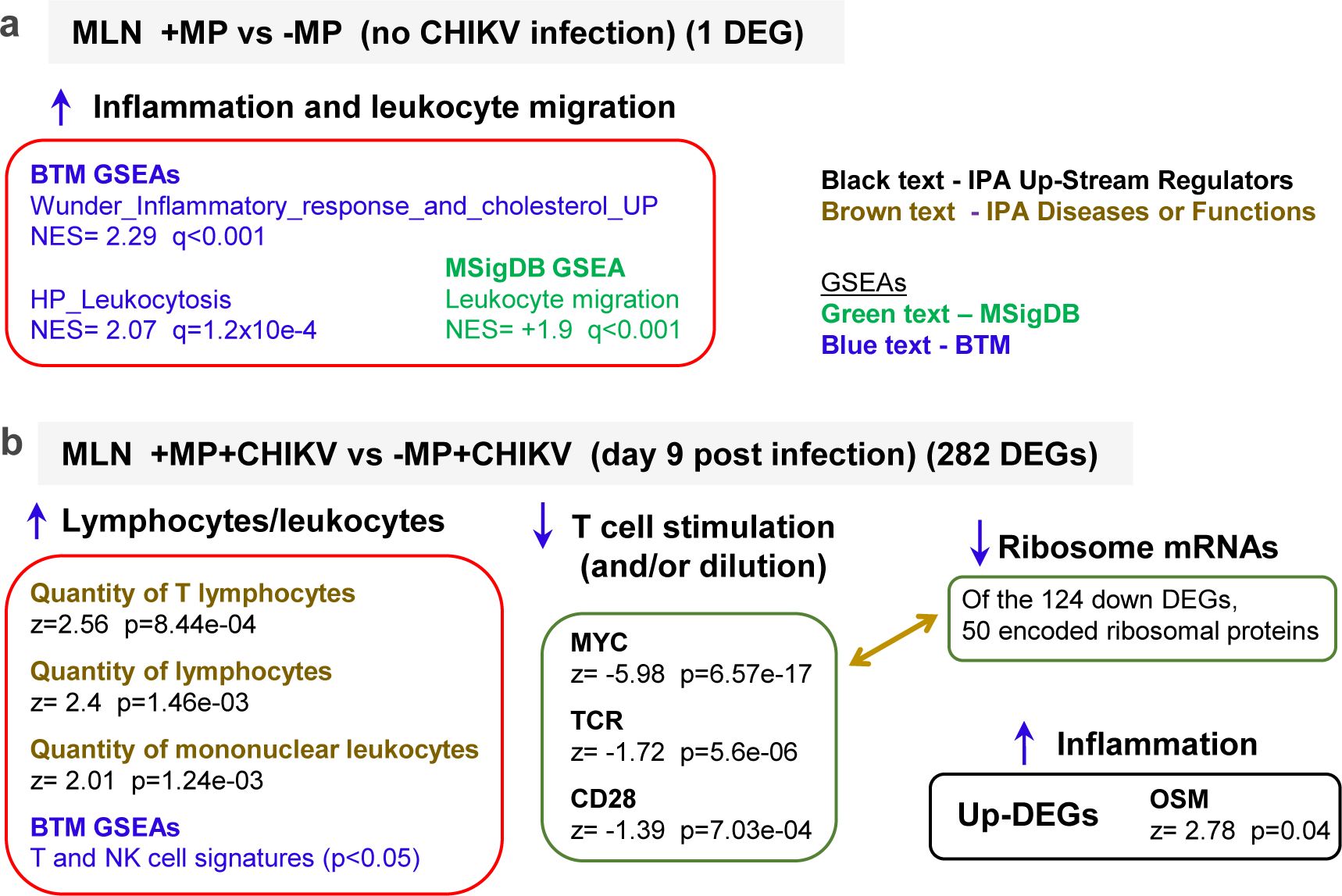
Analysis of RNA-Seq data from mesenteric lymph nodes. **a** RNA-Seq was used to compare mesenteric lymph nodes (MLNs) from mice drinking MP (+MP) with MLNs from mice drinking water without MP (-MP) for 28 days. The full gene list (Supplementary Table 4a) was interrogated using GSEAs and gene sets from MSigDB (Supplementary Table 4b) (green text) and BTMs (Supplementary Table 4c) (blue text). **b** RNA-Seq was used to compare MLNs 9 days after CHIKV infection for +MP+CHIKV vs –MP+CHIKV, with mice having their drinking water supplemented with or without MP for 37 days (i.e. 28 plus 9). DEGs (n=282) were identified (Supplementary Table 4f) and analysed by IPA (Supplementary Table 4g,h) and GSEAs (Supplementary Table 4i,j)

### MP consumption promoted lymphocytosis in MLNs 9 days after CHIKV infection

After 28 days +MP and -MP groups were infected with CHIKV, with RNA-Seq analysis of MLNs day 5 after CHIKV infection providing no compelling DEGs (Supplementary Table 4d). In contrast, on day 9 post infection 282 DEGs for +MP+CHIKV vs –MP+CHIKV were identified, although fold changes were low, ranging from log_2_ 0.87 to -0.71 (Supplementary Fig. 7b, Supplementary Table 4e,f). IPA analysis of the 282 DEGs provided a series of annotations indicating that the +MP+CHIKV MLNs had significantly more lymphocytes and T cells (Fig. 3b, Lymphocytes/leukocytes). In support of this finding, GSEAs using BTMs identified T cell and NK signatures in this group with significant positive NES scores (Fig. 3b; Supplementary Table 4i). Curiously, IPA USR analysis argued that T cell stimulation was decreased, with a highly significant negative z-score for MYC (Fig. 3b, T cell stimulation). MYC is a master regulator of metabolic programming in activated T cells (Shyer et al., 2020). TCR and CD28 (T cell costimulation) signatures also showed negative z-scores (Fig. 3b, T cell stimulation). The dominant feature of the DEG list was that 51 of the 124 down-regulated genes represented mRNAs encoding ribosomal proteins (Supplementary Table 4f) (Fig. 3b. Ribosome mRNAs).

Although many factors can regulate ribosome biogenesis (Petibon et al., 2021), a dominant feature of antigen-specific T cell activation is stimulation of ribosome biogenesis (Galloway et al., 2021). These results may suggest that, although there were more T cells in the MLN of the +MP+CHIKV group, overall they were less activated (Tan et al., 2017), with the lymphocytosis/leukocytosis perhaps diluting antigen-specific T cells.

As the down-regulated DEGs may be due (at least in part) to dilution effects, the up-regulated DEGs (n=158) were analyzed separately by IPA. This analysis thus reveals the processes that are being promoted, despite any dilution effects. OSM, a biomarker for inflammatory bowel disease (Verstockt et al., 2021), was returned as a significant USR (Fig. 3b, Inflammation; Supplementary Table 4g, column AC). OSM can also drive intestinal inflammation in mice (West et al., 2017) and can promote Th1 responses (Jung et al., 2010). Although overall fold changes were low, only one ribosomal protein was up-regulated, Rps6ka3 (RSK2) (Supplementary Table 4f), a protein associated with promoting Th17 responses in mice (Takada et al., 2016). Alerted CD4 T cell polarization were also suggested in the +MP+CHIKV group by GSEAs using MSigDB data sets (Supplementary Table 4j).

The IPA Diseases or Functions analysis returned a number of virus infection annotations with positive z scores, suggesting increased virus infection in day 9 MLNs from the +MP+CHIKV group, when compared with the -MP+CHIKV group (Supplementary Table 4h). These annotations are hard to reconcile with (i) the lack of differences in viremia (Fig. 2c, Supplementary Fig. 4a), (ii) although not significant, the lower viral loads in the colons of the +MP+CHIKV group (Supplementary Fig. 4d, e), and (iii) the lack of any viral reads in RNA-Seq data for day 9 MLN (Supplementary Fig. 7c). In addition, the up-regulated DEGs that gave rise to the virus infection annotations (Supplementary Fig. 7d) were all associated with negative regulation of metabolic processes when analyzed by Gene Ontology (Supplementary Fig. 7e).

This result is clearly not consistent with viral signatures, but is consistent with the aforementioned down-regulation/dilution of T cell activation signatures in +MP+CHIKV MLNs.

In summary, on day 9 post CHIKV infection, MP consumption was associated with a small change in the transcriptional profile in the MLNs that suggested a lymphocytosis/leukocytosis, with indications of increased inflammation and changes in T cell polarization.

### CHIKV arthritic foot swelling is significantly prolonged after MP consumption

After 28 days of drinking water supplemented with MP (+MP) or without MP (-MP), mice were infected with CHIKV. The food and water consumption was not significantly different for the +MP+CHIKV vs –MP+CHIKV groups after CHIKV infection (Supplementary Fig. 1a,c). The viremia was also not significantly different for these groups (Fig. 2c, Supplementary Fig 4b).

**Figure 4.**
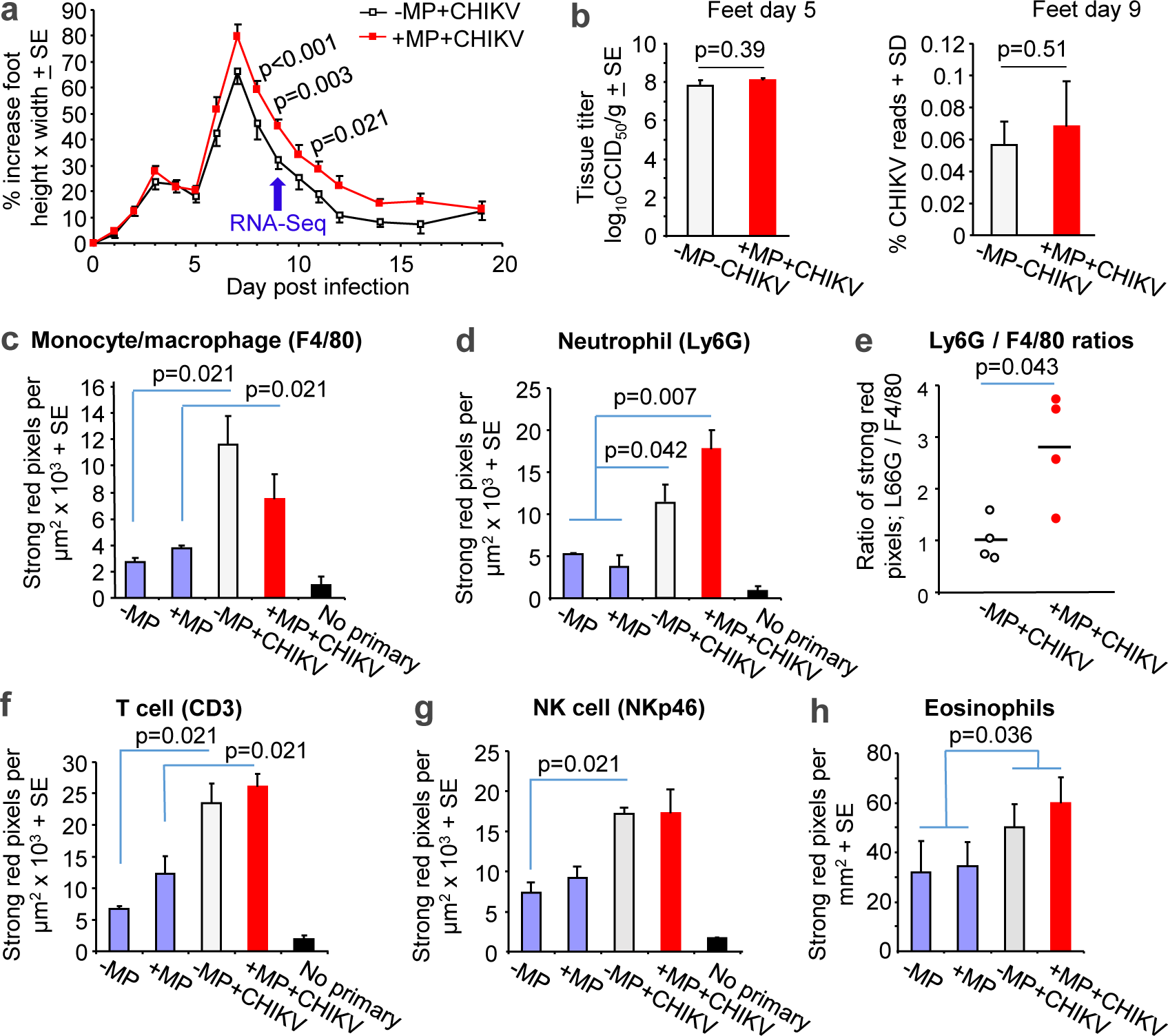
Infection and arthritis in +MP+CHIKV versus –MP+CHIKV. **a** Female C57BL/6J mice with drinking water supplemented with or without MP for 28 days were infected with CHIKV and foot swelling monitored. MP supplementation was continued until the end of the experiment. (n=24 feet from 12 mice day 0 to 5, n=16 feet from 8 mice from days 6 to 9, and n=8 feet from 4 mice from day 10 to19. Statistics by Mann Whitney U tests. Feet from 4 mice per group were harvested on day 9 for RNA-Seq (blue arrow). **b** The percentage of reads mapping to the CHIKV genome (n=4 feet from 4 mice per group) Statistics by t test. **c** IHC using an anti-F4/80 antibody. Statistics by Kruskal Wallis tests, n=4 feet from 4 mice per group, 3 sections per foot to produce a mean for each foot. **d** IHC using an anti-Ly6G antibody. Statistics as in c except n=2 feet for +MP and –MP groups, so these were combined for statistical purposes. **e** The ratio of Ly6G/ F4/80 staining for each foot from 4 mice per group. Statistics by Kruskal Wallis test. **f** IHC using an anti-CD3. Statistics as in c. **g** IHC using an anti-NKp46 antibody. Statistics as in c.

Overt CHIKV arthritic foot swelling in C57BL/6J mice usually peaks on day 6-7 post infection and then resolves (Gardner et al., 2010; Prow et al., 2019). Although the increased peak foot swelling between +MP+CHIKV and -MP+CHIKV groups did not reach significance, significantly greater foot swelling was seen on several days after the peak of foot swelling (Fig. 4a). The same result was seen in a repeat experiment, with significantly prolonged foot swelling again observed in the +MP+CHIKV group (Supplemental Fig. 8a). Feet from mice (n=4 feet from 4 mice per group) were taken day 9 post infection for RNA-Seq analysis (Fig. 4a, blue arrow), with these mice representing a subset of the mice shown in Fig. 4a. The swelling in these 4 feet per group was also significantly different for the +MP+CHIKV vs –MP+CHIKV groups (Supplemental Fig. 8b).

The prolonged foot swelling was not associated with increased viral loads in the feet as neither tissue titers on day 5, or CHIKV reads from RNA-Seq analysis of feet on day 9 post infection, showed any significant differences (Fig. 4b).

### MP consumption increases Ly6G/F4/80 ratios in arthritic feet after CHIKV infection

A prodigious inflammatory infiltrate is well described for CHIKV arthritis in this model (Gardner et al., 2010; Nguyen et al., 2020; Poo et al., 2014b; Wilson et al., 2017) and for alphaviral athritides generally (Lin et al., 2020; Suhrbier, 2019; Suhrbier et al., 2012). Although CHIKV infection (i.e. +MP+CHIKV vs +MP and -MP+CHIKV vs +MP) did show the expected significant increases, no significant differences in the level or density of leukocyte infiltrates was observed between +MP+CHIKV and –MP+CHIKV groups (Supplementary Fig. 8c,d). This argues that the increased foot swelling for the +MP+CHIKV vs MP+CHIKV group, was not predominantly due to increased levels of edema.

To investigate whether the composition of the inflammatory infiltrates were changed by MP consumption, immunohistochemistry (IHC) was undertaken on feet day 9 post infection (Supplementary Fig. 9) and staining quantitated by Aperio pixel count (Fig. 4c-g). F4/80 is generally considered to be a marker for tissue-residing monocytes (generally F4/80 low) and macrophages (F4/80 positive). A prodigious monocyte/macrophage infiltrate is well described for CHIKV arthritis and alphaviral athritides generally (Gardner et al., 2010; Lin et al., 2020; Suhrbier, 2019; Suhrbier et al., 2012). Thus as expected, IHC using a F4/80 showed significantly elevated staining after CHIKV infection. However, no significant difference (p=0.2, t test) between +MP+CHIKV and -MP+CHIKV groups was observed (Fig. 4c). Increased numbers of infiltrating neutrophils have been associated with exacerbation of arthritis in this model (Poo et al., 2014a; Prow et al., 2019) and other CHIKV mouse models (Cook et al., 2019). IHC using the neutrophil marker, Ly6G, showed significant increases associated with CHIKV infection (Fig. 4d), but comparison between +MP+CHIKV versus -MP+CHIKV did not reach significance (p=0.082, t test). However, when the Ly6G/F4/80 ratio was calculated for each foot, the +MP+CHIKV group showed a significant increase in this ratio (Fig. 4e). An increase in this ratio has been associated with a proinflammatory phenotype in a number of settings (Huang et al., 1999; Malerba et al., 2006; Martinez Gomez et al., 2013; Pawlaczyk et al., 2008).

T cells, and to a lesser extent NK cells, are known to promote arthritis in this model (Suhrbier, 2019; Teo et al., 2015). However here were no significant difference in CD3 (T cell) staining, between +MP+CHIKV and –MP+CHIKV groups, although as expected (Gardner et al., 2010), CHIKV infection induced a significant increase in CD3 staining (Fig. 4f). NK staining (NKp46) showed similar results to CD3 staining (Fig. 4g). Eosinophils were recently linked to skeletal muscle wound healing (Hazlewood et al., 2021) and may have an anti-inflammatory role in this CHIKV arthritis mouse model (Poo et al., 2014a). However, no significant differences in eosinophil staining was apparent between +MP+CHIKV and –MP+CHIKV groups (Fig. 4h; Supplementary Fig. 9).

### MP consumption has no discernable effect on the transcriptional profile in feet

Mice drinking MP-supplemented drinking water for 37 days (28 + 9 days) (no CHIKV infection) were analyzed by RNA-Seq (Supplemental Fig. 10a). Only one DEG, a putative gene, was identified for the +MP vs –MP comparison in feet before CHIKV infection (Supplementary Table 5a). The result is consistent with the lack of a discernable difference in foot measurements (Supplemental Fig. 10b) and IHC (Fig. 4c-h) for the +MP vs –MP groups in the absence of CHIKV infection.

### MP consumption prolongs Th1 responses in CHIKV arthritis

Mice drinking water with and without MP for 28 days were infected with CHIKV and on day 9 post infection feet were harvested for RNA-Seq. PCA plots showed a clear separation of the two groups (Supplementary Fig. 10c), with 1202 DEGs identified when a q<0.05 filter was applied (Supplementary Table 5b). As a large number of these DEGs had low fold changes, and in order to focus bioinformatic treatments on the more significant transcriptional changes, we increased the significance stringency to q<0.01, which provided 460 DEGs (Supplementary Table 5c).

Analyses of the 460 DEGs using a IPA, Cytoscape, and GSEAs with MSigDB and BTMs, provided multiple annotations associated with increased inflammation and immune activation for +MP+CHIKV vs -MP+CHIKV (Supplementary Table 5d-i), consistent with the increase foot swelling in the +MP+CHIKV group (Fig. 4a). Th1 CD4 T cells are thought to be the major drivers of CHIKV arthropathy (Suhrbier, 2019), with multiple signatures indicating increased Th1 bias in the arthritic feet of the +MP+CHIKV group (Fig. 5, Th1; Supplementary Table 5d,e,h). NK cells also contribute to arthritis in this model (Suhrbier, 2019; Teo et al., 2015), with the top IPA Canonical Pathway annotations indicating increased NK cell activity in +MP+CHIKV feet (Fig. 5, NK activity; Supplementary Table 5d). Some NK annotations were also seen in IPA Diseases or Functions (Fig. 5, NK activity; Supplementary Table 5f), and in GSEAs using BTMs (Supplementary Table 5i).

**Figure 5.**
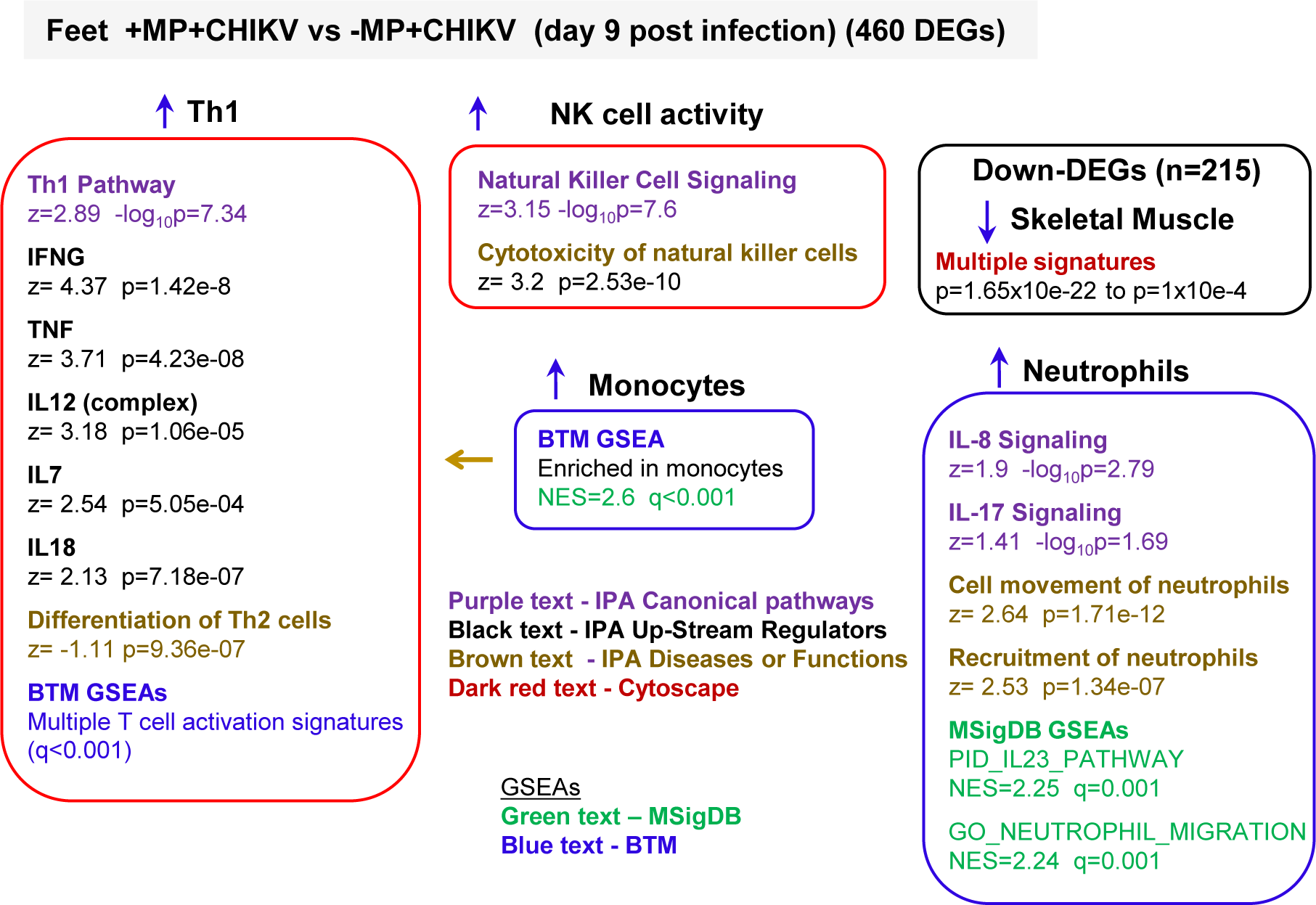
Analysis of RNA-Seq data from feet day 9 post infection for +MP+CHIKV versus–MP+CHIKV. RNA-Seq of feet, taken day 9 post infection, provided 460 DEGs (p<0.01) for +MP+CHIKV versus –MP+CHIKV (Supplementary Table 5b). These DEGs were analyzed by IPA and Cytoscape (Supplementary Table 5c-f) and the All gene list (Supplementary Table 5a), pre ranked by fold change, interrogated using GSEAs and gene lists from MSigDB and BTMs (Supplementary Table 5g,h). Top and other selected annotations are shown, highlighting the pathways known to be associated with promotion of arthritis.

Cytoscape analysis of the down-regulated DEGs was dominated by skeletal muscle annotations (Fig. 5, Skeletal Muscle; Supplemental Table 5g), indicating increased muscle cell damage in the +MP+CHIKV group. Myositis and muscle damage is well described in this mouse model (Gardner et al., 2010; Nakaya et al., 2012; Prow et al., 2019) and was clearly evident by H&E staining of feet taken day 9 post infection (Supplementary Fig 10d).

The BTM GSEAs suggested an increase in monocytes (Fig. 5, Monocytes; Supplementary Table 5h), perhaps consistent with the Th1 bias (De Koker et al., 2017; Iijima et al., 2011).

Macrophage annotations were ranked relatively lower, and suggested mild up-regulation of activity in the +MP+CHIKV group (Supplementary Table 5d,f,h). Although macrophages contribute to the pathogenesis of alphaviral arthritides, they are also important for disease resolution, which would be expected to be underway by day 9 (Prow et al., 2019). A transition from inflammatory macrophages to resolution phase macrophages (Schroder et al., 2019) is associated with removal of neutrophils, and represents a common theme for returning to hemostasis after tissue damage (Soehnlein and Lindbom, 2010). Using GSEA and a macrophage resolution signature described previously (Prow et al., 2019), a highly significant positive enrichment occurred in the +MP+CHIKV group, suggesting that increased inflammation led to increased resolution phase activity, and that prolonged foot swelling was not associated with a loss of resolution phase activity (Supplementary Fig. 10e).

Although relatively less dominant than the Th1 signatures, a series of neutrophil-associated annotations were evident in the +MP+CHIKV group (Fig. 5, Neutrophils; Supplementary Table 5d,f,h,i), consistent with the increased Ly6G/F4/80 staining ratios (Fig. 4e). Increased neutrophils have previously been associated with promotion of arthritis and foot swelling in this model (Poo et al., 2014a; Prow et al., 2019).

The IPA USR analysis of the 460 DEGs also provided a series of significantly up-regulated type I interferon response annotations (Supplementary Table 5e; Interferon alpha, IFNB1, IRF1, IRF7). Such annotations are prominently associated with CHIKV infection (Wilson et al., 2017); however, there were no significant differences in viral loads between +MP+CHIKV and – MP+CHIKV (Fig. 4d). The sustained IFNγ responses (Fig. 5, Th1) may have prolonged or sustained generation of type I IFN responses during the course of CHIKV infection (Eloranta et al., 1997) in +MP+CHIKV group.

### No significant effects of MP consumption on the gut microbiome

A number of studies in a range of species (Fackelmann and Sommer, 2019; Hirt and Body-Malapel, 2020), including mice (Jin et al., 2019; Li et al., 2020a; Lu et al., 2018), have described alterations in the gut microbiome or dysbiosis mediated by MP ingestion. In this CHIKV mouse model arthritis can also be influenced by a high fiber diet (Prow et al., 2019) and the microbiome can also influence antiviral resistance (Winkler et al., 2020). To determine whether the enhanced arthritis after MP consumption described above might be associated with MP-induced dysbiosis, the fecal microbiome was investigated over time using an experimental set-up that sought to minimize the influence of facility and cage effects on the results of the microbiome analyses (Basson et al., 2020; Parker et al., 2018). Such effects are now well recognized as potentially compromising interpretation of mouse microbiome studies (Basson et al., 2020; Parker et al., 2018). The results (Supplementary Table 6a-e) showed no significant differences for +MP versus -MP groups after 28 days of MP consumption at the genus (Fig. 6a), class (Supplementary Fig. 11) or phylum levels (Supplementary Fig. 12), with the comparisons involving 24 mice and 8 cages per group. Note both +MP and -MP had two independent replicate groups on days 0 and 28 (Fig. 6a).

**Figure 6.**
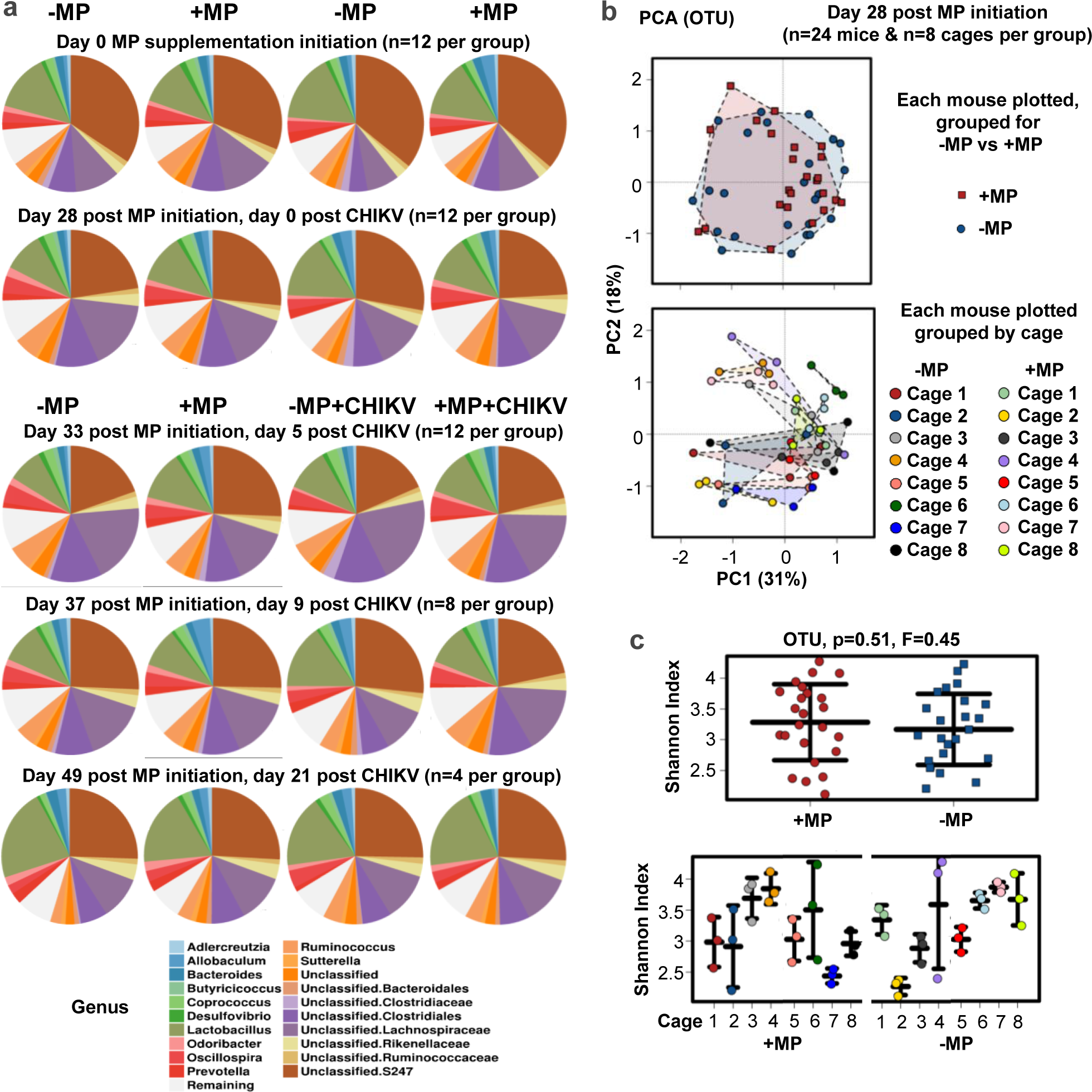
No effect of MP consumption on the microbiome. **a** Female C57BL/6J mice (n=48) were randomly assigned into four groups; each group comprised 12 mice housed in 4 cages. On day 0 two groups were given drinking water supplemented with MPs (+MP), with two groups left drinking unsupplemented water (-MP). On days 0 and 28 there were thus two replicate groups for +MP and two replicate groups for –MP (a total of n=24 mice and n=8 cages per group). On day 28, mice in one -MP and one +MP group were infected with CHIKV (- MP+CHIKV and +MP+CHIKV). On day 5 and 9 post CHIKV infection some mice were euthanized for histology and RNA-Seq. Fecal samples were taken from each (and each remaining) mouse at the indicated times and the microbiome determined by 16S sequencing and analyzed by Calypso, with the mean of all mice in each group shown. The genus levels output is shown. **b** On day 28 post onset of MP supplementation, all +MP versus all –MP mice were compared (n=24 mice and n=8 cages per group) by PCA analyses based on OTUs, with each plotted data point representing one mouse. The data is presented in two ways, +MP versus -MP (top) and by cage (bottom). **c** The Shannon indexes (based on OTUs) for all the mice in b plotted for +MP versus -MP (top) and by cage (bottom). Statistical comparison for +MP versus -MP by one way ANOVA, p=0.51, F-statistic=0.45.

Principle component analyses (PCA) (Fig. 6b, top graph) and Shannon Diversity Indices (Fig. 6c, top graph), both based on Operational Taxonomic Units (OTUs) (Mandal et al., 2015), again showed no significant effect on the microbiome after 28 days of MP consumption. As expected, individual cage effects (Basson et al., 2020) were clearly discernable (Fig. 6b and c; bottom graphs), but on average no significant effect of MPs consumption on the microbiome emerged.

### CHIKV infection can effect Firmicutes/Bacteroidetes ratios

On day 28 after initiation of MP consumption, two groups of mice were infected with CHIKV and the microbiome was analyzed on day 0, 5, 9 and 21 post infection (Supplementary Table 6a- i). After CHIKV infection, no significant differences emerged at any time point for +MP+CHIKV versus -MP+CHIKV (or the remaining +MP and -MP groups) at the genus (Fig. 6a), class (Supplementary Fig. 11) or phylum levels (Supplementary Fig. 12). However, from day 0 to day 5 in the -MP+CHIKV group, there was a significant increase in Firmicutes and a significant decrease in Bacteroidetes (Fig. 7a; Supplementary Table 6j). Although the +MP+CHIKV group had the same directions of change in these bacterial phyla, these did not reach significance (Fig. 7a; Supplementary Table 6j).

**Figure 7.**
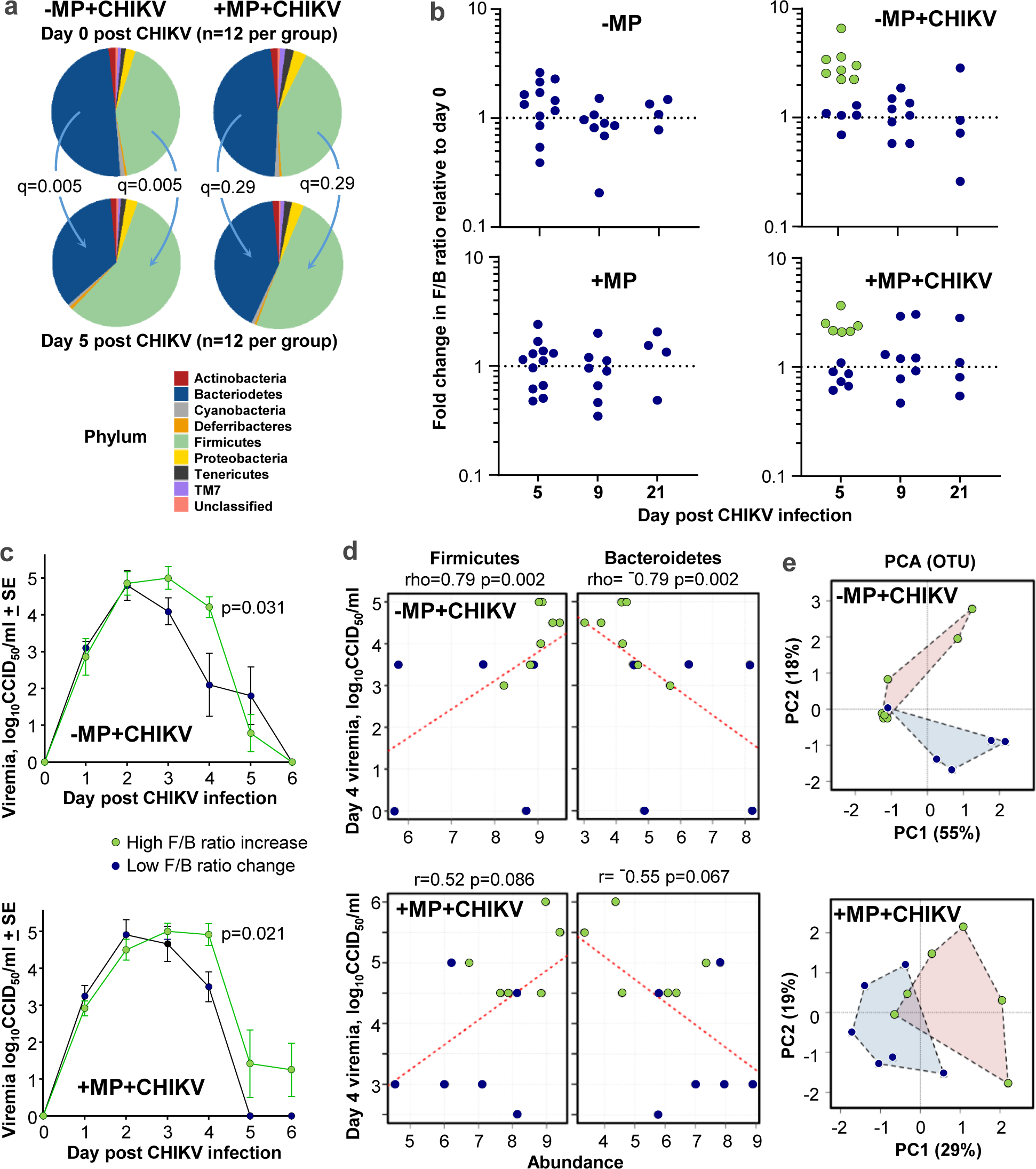
MP supplementation and the microbiome after CHIKV infection. **a** The microbiome analyzed by Calypso and plotted at the phylum level for mice day 0 and 5 post CHIKV infection. The microbiome from this subset of mice were previously described at the genus level in Fig. 6a. Changes in the proportion of Firmicutes and Bacteriodetes, which reach significance in the -MP+CHIKV mice, but not the +MP+CHIKV mice (Supplemental Table 6b). **b** The fold changes in the Firmicutes/Bacteriodetes (F/B) ratios in individual mice from day 0 to the indicated day post-CHIKV infection. In both -MP+CHIKV and +MP+CHIKV groups, on day 5 a subset of mice showed a relatively higher increase in F/B ratios when compared to day 0 (green circles), whereas the remaining mice showed a relatively lower change (blue circles). **c** The mean viremias for the mice that on day 5 had relatively high increases in F/B ratios compared with those with low change in F/B ratios. Same mice as shown in b for day 5. The differences in viremia were significant on day 4; -MP+CHIKV, statistics by Kruskal-Wallis test (non-parametric data distribution), and +MP+CHIKV by t test (parametric data distribution). **d** Correlations of fecal Firmicutes and Bacteriodetes abundance on day 5 versus viremia on day 4 for individual mice. Statistics, for -MP+CHIKV by Spearman correlations (non-parametric data distributions) for +MP+CHIKV by Pearson correlations (parametric data distributions). **e** PCAs illustrating that the groups identified as having a high increase (green circles) and a low change in F/B ratios (blue circles) have microbiomes that also clearly segregate at the OTU level on day 5.

Changes in Firmicutes/Bacteroidetes (F/B) ratios are widely accepted as representing a biomarker of dysbiosis (Stojanov et al., 2020), so the fold changes in F/B ratios from day 0 to days 5 , 9 and 21 were plotted for individual mice (Fig. 7b). On day 5 post infection, mice appeared to segregate into two populations, with 8/12 mice in the -MP+CHIKV group and 6/12 in the +MP+CHIKV group showing a relatively high increase in their fecal F/B ratios (Fig. 7b, green data points), compared to the remaining mice that showed only a low change (Fig. 7b, blue data points). When the viremias for the two populations where plotted they showed a significant difference on day 4 post CHIKV infection for both the -MP+CHIKV and the +MP+CHIKV groups (Fig. 7c). However, increased or low F/B ratio changes and the associated effects on viremia on day 4, did not translate to significant differences in foot swelling (Supplementary Fig. 13). Increased viremia on day 4 also correlated with increased abundance of Firmicutes and decreased abundance of Bacteroidetes, although in the +MP+CHIKV group this only approached significance (Fig. 7d). Finally, the distinction between high and low F/B ratio changes (at the phylum level) was maintained at the OTU level, with PCA plots showing a clear separation (Fig. 7e).

There was therefore no indication from these studies that MP consumption had a significant effect on the microbiome. Nevertheless, CHIKV infection did appear to affect the microbiome, with increased viremias associated with relatively high increases in F/B ratios. Increases in Firmicutes were primarily driven by increases at the genus level of Lachnospiraceae and Clostridiales, and decreases in Bacteroidetes primarily driven by a decrease at the genus level of S247 (Supplementary Table 6k). That a higher F/B ratio caused the increases in viraemia is perhaps less likely given (i) the F/B ratio change was relative to day 0 post infection and (ii) a Clostridium species (phylum Firmicutes) was shown to promote antiviral activity (Winkler et al., 2020), whereas in our data more Firmicutes correlated with more virus (Fig. 7d).

## DISCUSSION

We show herein that the consumption of MP predisposes mice to a prolonged arthritis after CHIKV infection. This occurred in the absence of any detectable MPs entering the body or any significant effects of MP consumption on the microbiome. RNA-Seq and bioinformatic analyses argue that MP consumption leads to a mild increase in gut inflammation. This then predisposed the viral arthritis to an extended immunopathology characterized by elevate Th1 biased, and to a lesser extent, Th17 biased, signatures.

The picture that emerges is reminiscent of enteropathic arthritis. A connection between bowel pathology and arthritis has long been recognized (Ashrafi et al., 2021), with the arthritis that is often associated with Inflammatory Bowel Disease (IBD) now referred to as enteropathic arthritis (Farisogullari et al., 2021; Gutiérrez-Gonzalez et al., 2020; Picchianti-Diamanti et al., 2020). The mechanism(s) underpinning the causal relationship whereby inflammation in the gut leads to inflammation in the joints remain poorly characterized and often speculative.

Nevertheless, migration of Innate Lymphoid Cells (ILCs), in particular ILC1 and ILC3 have recently been implicated, with these cells detecting gut damage or perturbation, becoming activated and migrating into joint tissues and promoting Th1 and Th17 driven arthritis (Fang et al., 2020; Neerinckx et al., 2015). ILCs are broadly divided into three categories ILC1, ILC2 and LC3 cells, which to some extent mirror the function of Th1, Th2 and Th17 T cells, respectively, without displaying any antigen specificity (Jarade et al., 2021; Vivier et al., 2018). We derived gene sets from a series of papers and used these in GSEAs to identify potential ILC signatures (Supplementary Table 7). After CHIKV infection, MP consumption was associated with significantly enriched ILC1 signatures in colon, MLN and feet, with ILC3 signatures seen in MLNs and feet, and ILC2 signatures seen only in feet (Supplementary Fig. 14). These analyses are consistent with the proposed model of enteropathic arthritis (Fang et al., 2020), although ILC1 signatures would be difficult to distinguish from NK signatures (McFarland et al., 2021) in whole tissues where NK cells are also present (Fig. 5, NK cell activity). Another role of ILCs is protection of the gut from infection (Shannon et al., 2021) and maintenance of gut homeostasis (Diefenbach et al., 2020), which may explain the slightly lower viral loads in the colon (Supplementary Fig. 4d,e), lower epithelial repair signatures in the colon (Fig. 2d) and the better weight recovery in the +MP+CHIKV group (Fig. 4a). Faster weight recovery and prolonged arthritis after CHIKV infection did not correlate (Supplementary Fig. 15), supporting the notion that these are independent outcomes of MP consumption and ILC stimulation.

A central question remains regarding how low levels of MP consumption might cause gut perturbations. One might speculate this involves low level (Supplementary Table 1b), histologically undetectable (Supplementary Fig. 5a), epithelial cell plasma membrane disruption in the colon. This contention is consistent with (i) the signatures described in Fig. 2b, which suggest a plasma membrane signaling response (Le Roux et al., 2019; Morigasaki et al., 2019), (ii) the lack of MP uptake (Fig. 1b), and (iii) the large volume of literature on plasma membrane damage by MPs in cell lines *in vitro,* which also report oxidative stress as a consequence (Banerjee and Shelver, 2021; Wang et al., 2021a). Although MPs are well recognized as ideal abrasive agents for cosmetics (Bilal et al., 2020; Kozlowska et al., 2019), most humans and animals constantly swallow sand dust, which might be expected to have similar abrasive effects (Karadima et al., 2021). One could speculate that specific hydrophobicity characteristics of MPs (Pham et al., 2021) may facilitate their ability to perturb or disrupt the plasma membrane directly (McKarns et al., 1997; Ogbourne et al., 2004). Alternatively, although we were unable to see any significant overall differences in the microbiota, MPs coated with certain bacteria/biofilms (Miao et al., 2019; Ogonowski et al., 2018; Ribet and Cossart, 2015; Wu et al., 2019) may somehow adversely affect enterocyte plasma membranes (Ammendolia et al., 2021; Peterson and Artis, 2014). Another conceivable possibility is that MPs adsorb (Yu et al., 2021) certain chemicals (e.g. hydrophobic bile salts (Sagawa et al., 1993)) that that then imbues the MP with pathogenic properties (Stenman et al., 2013). Future mechanistic studies are clearly warranted.

Although the mechanism of gut perturbation remains speculative, this paper provides compelling evidence that MP consumption, without causing overt microbiome or gut changes or entering the body, can nevertheless significantly exacerbate the immunopathology associated with a viral arthritis. If MPs do indeed activate ILCs in the gut, and ILCs become a systemic immunological conduit (Fang et al., 2020; Neerinckx et al., 2015), one might speculate that MPs could dysregulate inflammation in a range of conditions (Benezech and Jackson-Jones, 2019; Fernando et al., 2021; Maleki et al., 2020).

## Acknowledgements

From QIMR Berghofer MRI we thank Dr I Anraku for managing the PC3 (BSL3) facility, Dr Clay Winterford and Crystal Chang for the histology and immunohistochemistry, the animal house staff for mouse breeding and agistment, and Dr. Gunter Hartel for assistance with statistics. We also thank Dr S.M. Bengtson Nash (Griffith University) for helpful discussions.

## Author Contributions

Conceptualization, A.S.; Methodology, D.J.R. and A.S.; Formal analysis, D.J.R., A.S., T.D., and C.B.; Investigation, D.J.R., T.L., K.Y, and B.T.; Data curation, D.J.R., A.S., T.D., and C.B.; Writing – original draft, A.S. and D.J.R.; Writing – review and editing, A.S., D.J.R. and T.D.; Visualization, D.J.R., A.S., T.D., and C.B.; Supervision, D.J.R. and A.S.; Project administration, A.S.; Funding acquisition, A.S.

## Declaration of Interests

The authors declare no competing interests.

## Funding

The work was funded by the National Health and Medical Research Council (NHMRC) of Australia (Investigator grant APP1173880 awarded to A.S.). The funders had no role in study design, data collection and analysis, decision to publish, or preparation of the manuscript.

## RESOURCE AVAILABILITY

### Lead contact

Further information and requests for resources and reagents should be directed to the Lead Contact, Andreas Suhrbier.

### Materials availability

Materials generated in this study will be made available on request, but we may require a completed materials transfer agreement.

### Data and code availability

All data is provided in the manuscript and accompanying supplementary files. Raw sequencing data (fastq files) generated for this publication for RNA-Seq and for the microbiome has been deposited in the NCBI SRA, BioProject: PRJNA753548 and are publicly available as of the date of publication.

## EXPERIMENTAL MODEL AND SUBJECT DETAILS

### Ethics Statement and PC3/BSL3 certifications

All mouse work was conducted in accordance with the “Australian code for the care and use of animals for scientific purposes” as defined by the National Health and Medical Research Council of Australia. Mouse work was approved by the QIMR Berghofer Medical Research Institute animal ethics committee (P2235 A1606-618M), with infectious CHIKV work conducted in a biosafety level3 (PC3) facility at the QIMR Berghofer MRI (Australian Department of Agriculture, Water and the Environment certification Q2326 and Office of the Gene Technology Regulator certification 3445).

### Mice and CHIKV infection

Female C57BL/6J mice (6-8 weeks old) were purchased from Animal Resources Center (Canning Vale, WA, Australia). CHIKV infection was undertaken as described (Gardner et al., 2010; Nguyen et al., 2020; Poo et al., 2014b) with 10^4^ CCID_50_ C6/36-derived CHIKV (Reunion Island isolate, LR2006-OPY1, GenBank: KT449801.1) inoculated subcutaneously (s.c.) into each hind foot. Virus was checked for mycoplasma as described (La Linn et al., 1995).

Swelling of hind feet was measured using digital calipers and is presented as a group average of the percentage increase in foot height × width for each foot compared with the same foot on day 0 (Nguyen et al., 2020). Serum viremia was determined using CCID_50_ assays of serum obtained by tail bleed as described (Gardner et al., 2010; Nguyen et al., 2020; Poo et al., 2014b). Intestine and feet tissue titers were determined by CCID_50_ assays of supernatants of homogenized tissues as described (Gardner et al., 2010). Tissues were also placed into RNAprotect Tissue Reagent (QIAGEN) for RT-qPCR and/or RNA-Seq analyses.

### Cell lines

Vero cells (ATCC#: CCL-81) and C6/36 cells (ATCC# CRL-1660) cell were maintained in RPMI 1640 (Thermo Fisher Scientific, Scoresby, VIC, Australia) supplemented with endotoxin free 10% heat-inactivated fetal bovine serum (FBS; Sigma-Aldrich, Castle Hill, NSW, Australia) at 37°C and 5% CO_2_. Cells were checked for mycoplasma using MycoAlert Mycoplasma Detection Kit (Lonza, Basel, Switzerland). FBS was checked for endotoxin contamination before purchase as described (Johnson et al., 2005).

## METHOD DETAILS

### Water supplementation with MPs

The MPs comprised Fluoresbrite® yellow-green polystyrene-based microspheres with a diameter of 1.00 µm (Cat# 17154-10) purchased from Polysciences as 2.5% w/v in aqueous suspension without sodium azide. The 1 µm microspheres were washed 3 times in 1 ml drinking water by centrifugation at 12,000 × g for 10 min, with the final wash including a 1 hr incubation on a rotor at 4°C. The MPs were diluted in mice drinking water to 10^6^ microspheres/ml (526 µg/l). Drinking water bottles containing MPs were inverted every 2-3 days, and fresh water with/without MP was provided every week. Mice were exposed to MP in their drinking water for 4 weeks prior to CHIKV infection, and thereafter until euthanasia.

For the water preference experiment, mice cages were fitted with 2 bottles of drinking water; one with and one without 1 µm MPs. Water in drinking bottles was measured using standard scales.

### MP detection in tissues and feces

Tissue and feces were weighted, manually chopped using scissors and digested in ammonium sulphate (50 mM), SDS (5 mg/ml) and proteinase K (1 mg/ml) overnight at 37°C as described (Walczak et al., 2015). Digested tissue suspensions were viewed by florescent microscope and a haemocytometer, with florescent MPs counted manually.

### CCID_50_ assays

Serum CCID_50_ assays were conducted as described previously using 10 fold serial dilution in duplicate in C6/36 cells and virus followed by detection using cytopathic effects in Vero cells (Gardner et al., 2010; Nguyen et al., 2020). For tissue titer determinations, tissues were weighed and placed in tubes containing RPMI1640 supplemented with 2% FCS and 4 x 2.8 mm ceramic beads (MO BIO Inc., Carlsbad, USA). Tissues were homogenized using Precellys24 Tissue Homogeniser (Bertin Technologies, Montigny-le-Bretonneux, France) 6000 x g for 15 seconds. After centrifugation twice at 21000 x g for 5 min titers in the supernatants were determined by CCID_50_ assays (as above) using 5 or 10 fold serial dilutions. The virus titers were determined by the method of Spearman and Karber.

### RT-qPCR

Mice feet or intestine was transferred from RNAprotect Tissue Reagent (QIAGEN) to TRIzol (Life Technologies) and was homogenized as above twice at 6000 x g for 15 sec. Homogenates were centrifuged at 14,000 × g for 10 min and RNA was isolated as per manufacturer’s instructions. cDNA was synthesized using iScript cDNA Synthesis Kit (Bio-Rad) and qPCR performed using iTaq Universal SYBR Green Supermix (Bio-Rad) as per manufacturer instructions with the following primers; CHIKV E1 (Poo et al., 2014b) Forward 5’- AGCTCCGCGTCCTTTACC -3’ and Reverse 5’- CAAATTGTCCTGGTCTTCCTG -3’, mRPL13a (Schroder et al., 2010) Forward 5’- GAGGTCGGGTGGAAGTACCA -3’ and Reverse 5’- TGCATCTTGGCCTTTTCCTT -3’. Serially diluted sample was used to generate a standard curve for CHIKV and mRPL13a primers, and this was used to calculate a relative quantity. qPCR reactions were performed in duplicate and averaged to determine the relative quantity in each sample, and CHIKV was normalized to mRPL13a to give relative CHIKV RNA levels.

### RNA-Seq and differential expression analyses

TRIzol extracted RNA from mice feet, large intestine and mesenteric lymph nodes was treated with DNase (RNase-Free DNAse Set (Qiagen)) followed by purification using RNeasy MinElute Cleanup Kit (QIAGEN) as per manufacturer instructions. RNA concentration and quality was measured using TapeStation D1K TapeScreen assay (Agilent). cDNA libraries were prepared using the Illumina TruSeq Stranded mRNA library prep kit and the sequencing performed on the Illumina Nextseq 550 platform generating 75 bp paired end reads. Per base sequence quality for >90% bases was above Q30 for all samples.

The quality of raw sequencing reads was assessed using FastQC (Simons, 2010)(v0.11.8) and trimmed using Cutadapt (Martin, 2011) (v2.3) to remove adapter sequences and low-quality bases. Trimmed reads were aligned using STAR (Dobin et al., 2013) (v2.7.1a) to a combined reference that included the mouse GRCm38 primary assembly and the GENCODE M23 gene model (Harrow et al., 2012) and CHIKV (GenBank KT449801.1, 11796 bp). Mouse gene expression was estimated using RSEM (Li and Dewey, 2011) (v1.3.0). Reads aligned to CHIKV were counted using SAMtools (Li et al., 2009) (v1.9). Differential gene expression in the mouse was analyzed using DESeq2 (v1.32.0) (Love et al., 2014) with default settings in R (v4.1.0), which employs Independent Filtering to remove genes with low expression values (Team, 2013).

Principle components analyses of log_2_ normalized counts were performed using ggpubr (v0.4.0) in R. For the 281 differentially expressed colon genes, expression values were compared by plotting the log_2_ ratio of normalized counts to row means in a heatmap that used Euclidean distance to cluster genes and samples, with the pheatmap package (v1.0.12) (Kolde, 2015) in R.

### Pathway Analyses

Up-Stream Regulators (USR), Diseases or Functions and Canonical pathways enriched in differentially expressed genes in direct and indirect interactions were investigated using Ingenuity Pathway Analysis (IPA) (QIAGEN).

### Functional Enrichment Analyses

Enrichment for biological processes, molecular functions, KEGG pathways and other Gene Ontology categories in DEG lists was undertaken using the STRING functional enrichment analysis in Cytoscape (v3.7.2)(Shannon et al., 2003; Szklarczyk et al., 2019). Gene Ontology (Ashburner et al., 2000; Gene Ontology, 2021) was also used to interrogate gene lists.

### Gene Set Enrichment Analyses

Preranked GSEA (Subramanian et al., 2005) was performed on a desktop application (GSEA v4.0.3) (Broad Institute, UCSanDiago) using the “GSEAPreranked” module. Differentially expressed “All gene” lists (after independent filtering in DESeq2), ranked by fold change, were interrogated for enrichment of gene sets from the complete Molecular Signatures Database (MSigDB) v7.2 gene set collection (31,120 gene sets) (msigdb.v7.2.symbols.gmt) and from Blood Transcription modules (BTMs)(Li et al., 2014).

We constructed gene sets for Innate Lymphoid Cells ILC1, ILC2 and ILC3 (Supplementary Table 7a) using a comprehensive series of references (Fang et al., 2020; Klose and Artis, 2020; McFarland et al., 2021; Panda and Colonna, 2019) and used these gene sets in GSEAs.

### Histology and immunohistochemistry

Feet histology, immunohistochemistry and quantitation of stain were undertaken as previously described (Gardner et al., 2010; Poo et al., 2014a; Prow et al., 2019; Wilson et al., 2017).

Briefly, feet were fixed in 10% formalin, decalcified with EDTA, embedded in paraffin and sections stained with hematoxylin and eosin (H & E; Sigma-Aldrich, Darmstadt, Germany). For immunohistochemistry, sections were stained with rabbit anti-CD3 (A0452; Dako, North Sydney, Australia), rat anti-F4/80 (ab6640; Abcam, Melbourne, Australia), goat anti-NKp46 (R&D Systems, AF2225), or rat anti-Ly6G (ab2557 NMP-R14; Abcam, Cambridge, MA, USA). Signal was detected using MACH2 Rabbit alkaline phosphatase (for CD3), Rat on Mouse alkaline phosphatase (for F4/80 & Ly6G) or Goat on Rodent alkaline phosphatase (for NKp46) Polymer Detection kits with Warp Red Chromogen (alkaline phosphatase) (Biocare Medical, Pacheco, CA, USA) (for CD3, F4/80 and NKp46), or NovaRed Chromogen Kit (peroxidase) (Vector Laboratories, Burlingame, CA, USA) (for Ly6G). Slides were scanned using Aperio AT Turbo (Aperio, Vista, CA USA) and analyzed using Aperio ImageScope software (Leica Biosystems, Mt Waverley, Australia) (v10) and the Positive Pixel Count v9 algorithm.

Eosinophils were labelled with Akoya Opal 620 tyramide (Akoya Biosciences, Marlborough, MA, USA) (Acharya and Ackerman, 2014; Liu et al., 2006) and staining analyzed by Aperio FL slide scanner (Leica Biosystems) and Qupath v0.2.3. (Bankhead et al., 2017).

For colon, tissue was fixed in 10% formalin, embedded in paraffin and stained with Alcian blue pH 2.5/periodic acid-Schiff (AB/PAS) stain. Quantitation of cells stained dark blue/purple (neutral mucins in goblet cells) was undertaken using Aperio Pixel count.

### 16S RNA-Seq of mice feces

Female C57BL/6J mice (n=48) were randomly placed in holding cages at QIMR Berghofer MRI animal facility (n=6 mice per cage) for 1 week and were fed standard chow and water (Parker et al., 2018). Mice were then reassigned to new experimental cages so that each experimental cage contained 3 mice (i.e. 50% of the standard mouse density of 6 mice per cage (Basson et al., 2020)), with each mouse taken from a different holding cage. After another week on the same chow and water, the cages were randomly allocated into four groups, each group comprising 12 mice in 4 cages. Two groups were then started on MP-supplemented water. This strategy was adopted to minimize the influence of cage effects on the results of the microbiome analysis (Basson et al., 2020).

Fecal pellets were collected fresh from each mouse in a cage directly into a screw cap micro tube (Sarstedt) at the indicated times. Pellets were heated at 100°C for 5 min, and sent to Australian Genome Research Facility (AGRF) for RNA extraction and 16S sequencing using V3-V4 region primers (Forward 5’- CCTAYGGGRBGCASCAG -3’ and Reverse 5’- GGACTACNNGGGTATCTAAT -3’. Sequencing was performed on an Illumina MiSeq platform.

### 16S bioinformatics

The bioinformatics analysis involved demultiplexing, quality control, OTU clustering, and taxonomic classification. Paired-ends reads were assembled by aligning the forward and reverse reads using PEAR (version 0.9.5). Primers were identified and trimmed. Trimmed sequences were processed using Quantitative Insights into Microbial Ecology (QIIME 1.8.4), USEARCH (version 8.0.1623), and UPARSE software. Using USEARCH tools, sequences were quality filtered, full length duplicate sequences were removed and sorted by abundance. Singletons or unique reads in the data set were discarded. Sequences were clustered and chimeric sequences were filtered using the “rdp_gold” database as a reference. To obtain the number of reads in each OTU, reads were mapped back to OTUs with a minimum identity of 97%. Taxonomy was assigned using QIIME. All samples passed quality control metrics.

Multivariate statistical analysis of the 16S data was performed using Calypso software (v 8.84) (Zakrzewski et al., 2017) with a .biom mapping file and metadata files. Data was filtered by excluding taxa that had less than 0.01% relative abundance, followed by total sum normalization (TSS) combined with square root transformation (Hellinger transformation).

## QUANTIFICATION AND STATISTICAL ANALYSIS

Statistical analyses of experimental data were performed using IBM SPSS Statistics for Windows, Version 19.0 (IBM Corp., Armonk, NY, USA). The t-test was used when the difference in variances was <4, skewness was >-2 and kurtosis was <2. Otherwise, the non-parametric Kolmogorov-Smirnov or Kruskal-Wallis tests were used.

## Supplementary Material

The Supplementary Material for this article comprises Supplementary Figures (pdf) and Supplementary Tables (xlsx).

## Notes

https://dataview.ncbi.nlm.nih.gov/object/PRJNA753548?reviewer=v1nkckn2fm0i69831e9989u1l7

### Competing Interest Statement

The authors have declared no competing interest.

