## Supplementary figures for "Microplastic consumption induces inflammatory signatures in the colon and prolongs a viral arthritis"

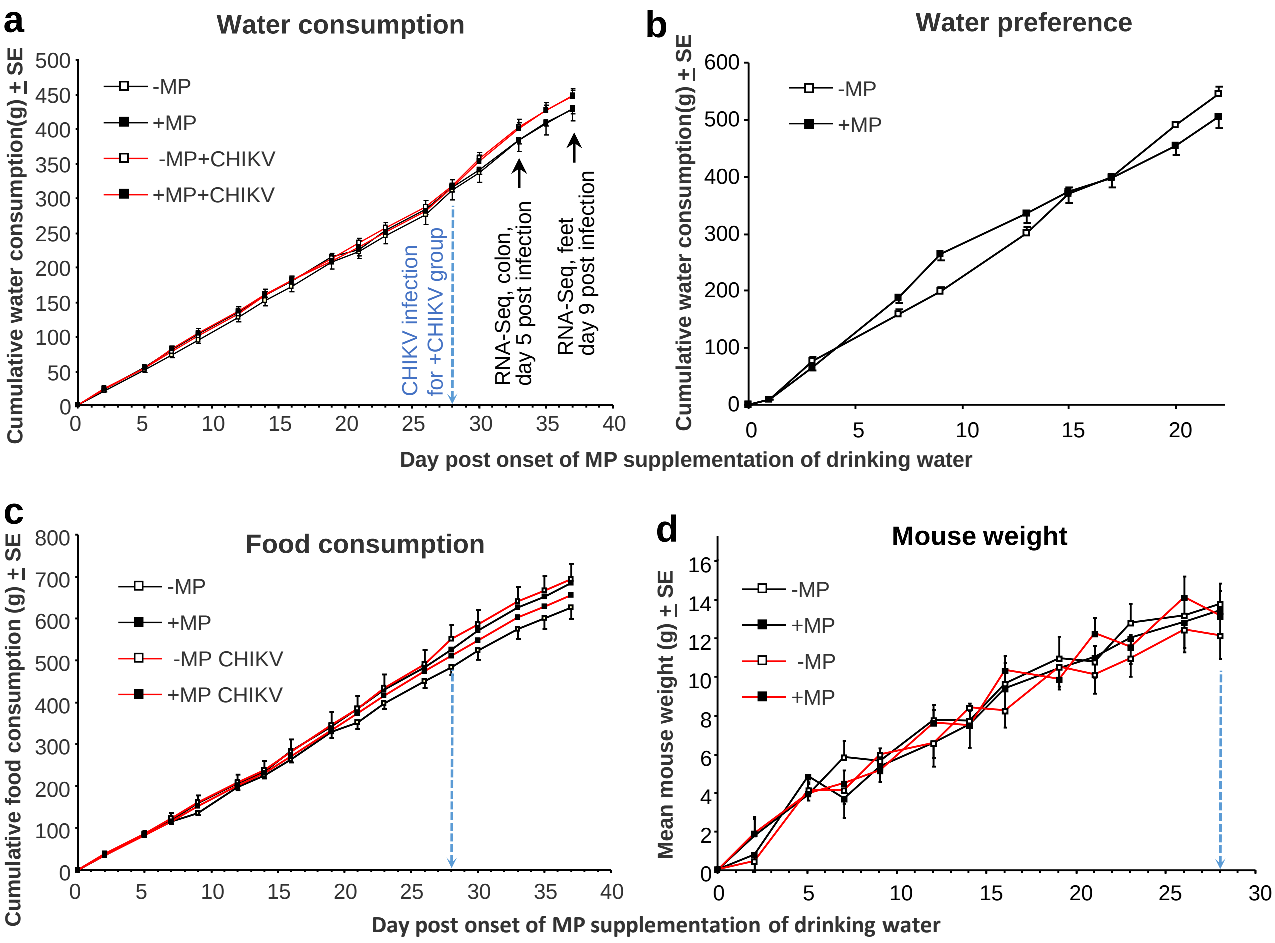


**Supplementary Fig. 1. Water and food consumption and body weight.** **a** Cumulative water consumption. Mean per cage for four cages each containing 3 mice; each group thus contained 12 mice. Half the mice were infected on day 28 with CHIKV, and on day 5 and 9 after CHIKV infection, 4 mice per group were euthanized for RNA Seq. Colons; the +MP and –MP groups had been drinking water with and without MP for 33 days, when colons were harvested. The +MP+CHIKV and –MP+CHIKV groups had been drinking water with and without MP for 33 days when colons were harvested and this time point also represented 5 days post infection. Feet; the +MP and –MP groups had been drinking water with and without MP for 37 days, when feet were harvested. The +MP+CHIKV and –MP+CHIKV groups had been drinking water with and without MP for 37 days when feet were harvested and this time point also represented 9 days post infection. **b** Water preference. In a separate experiment groups of mice were provided 2 water bottles, one with MP one without MP in each cage. 6 mice per cage, mean of 3 cages is shown. No significant differences emerged, illustrating that mice did not show a preference for MP-free water. **c** Cumulative food consumption for mice described in a. Mean per group is shown, with each group comprising four cages each containing 3 mice. **d** Mean mouse weight over time for mice in a, prior to CHIKV infection. n=12 mice per group. For weight change after CHIKV infection see Supplemental Fig. 4a.

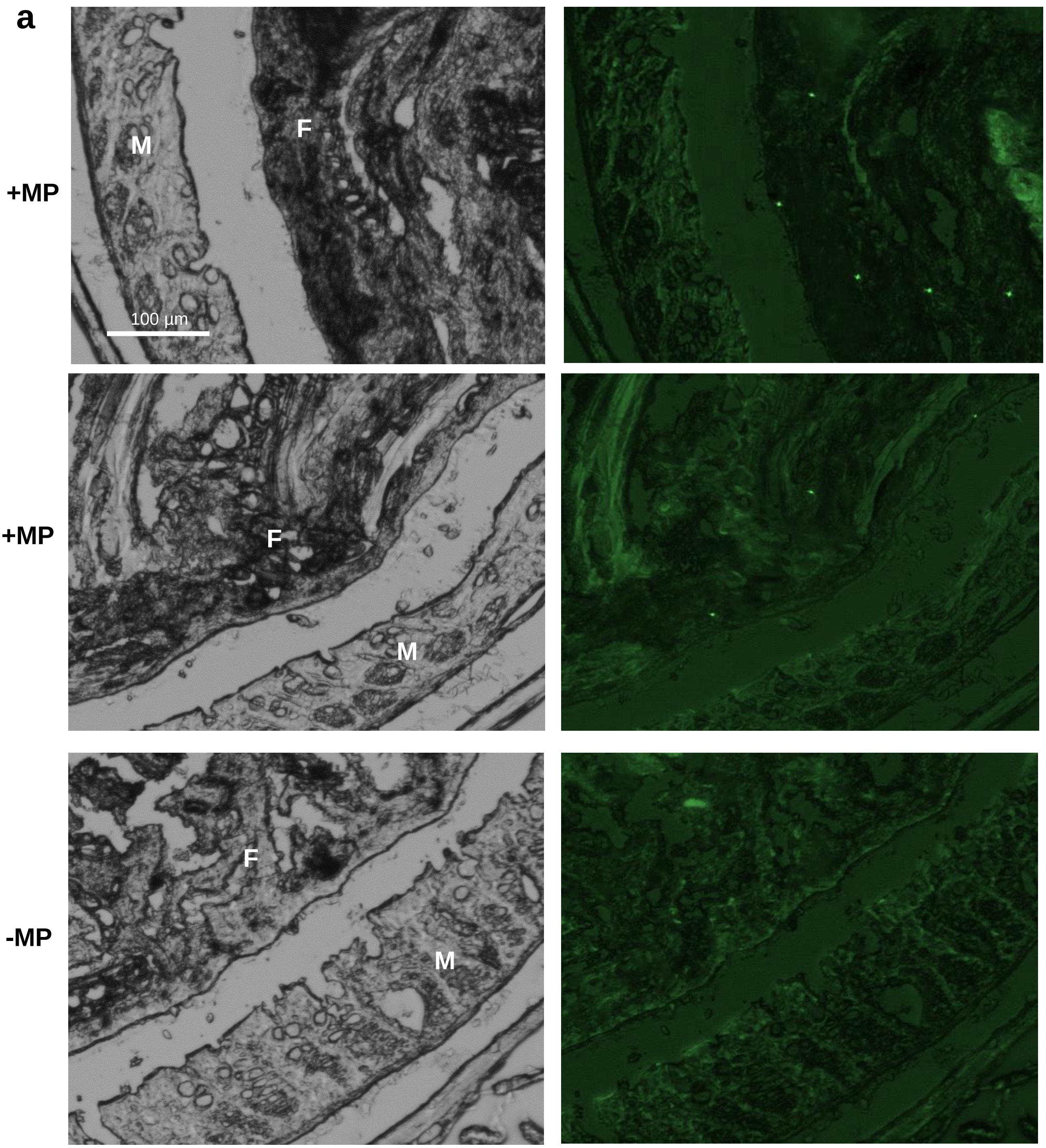


**Supplementary Fig. 2. MPs in faeces in the colon.** **a** Frozen sections of paraformaldehyde fixed colon from mice whose drinking water was (+MP) or was not (-MP) supplemented with MP for 8 weeks. Phase images left show colonic mucosa (M) adjacent to faecal material (F). Fluorescent images of the same sections (right) show the presence of MP in the faecal material but not mucosa for +MP mice. No MP were detected in the –MP group. Autoflurescence of faecal matter is dim and/or amorphous, readily distinguishable from the bright, discrete punctate fluorescent MPs.

### Colon (+MP vs –MP)

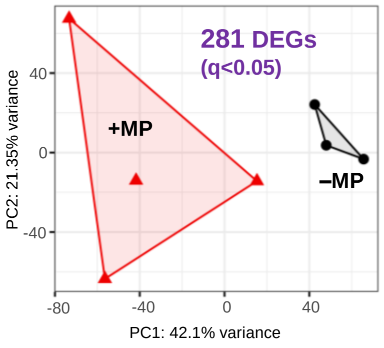


**Supplementary Fig. 3. PCA plot for RNA-Seq data from colon.** PCA plot for +MP versus –MP for the first two principal components of the log2 normalised counts of all genes that passed independent filtering by DESeq2. Derived from RNA-Seq data from colon taken from mice after 33 days of consumption of water with (+MP) and without MPs (-MP).

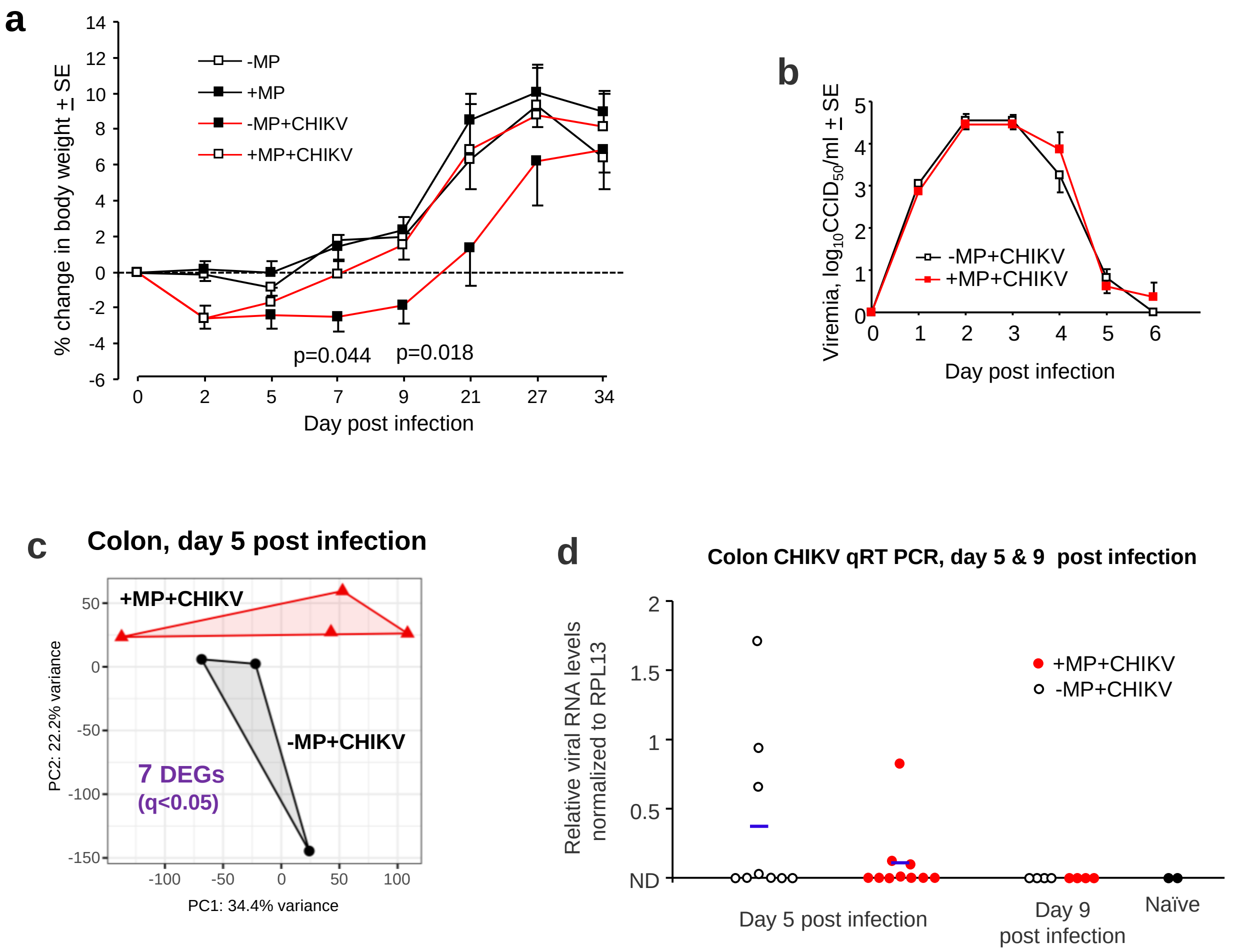


**Supplementary Fig. 4. Weight, viraemia, PCA and colon viral loads after CHIKV infection.**
**a** Weight change for the mice shown in Fig. 4a. Statistics by t tests; n=8 mice per group until day 9, thereafter n=4 mice per group. **b** Repeats of experiment shown in Fig. 2c. Mean viremia for a total of 14 mice per group from 2 independent experiments (that are also independent of the mice shown in Fig. 2c). **c** PCA plot of RNA-Seq data (for gene lists see Supplementary Table 2) for the first two principal components of the log2 normalised counts of all genes that passed independent filtering by DESeq2. **d** CHIKV qRT PCR of colons. Data for day 5 comes from 2 independent experiments; blue lines represent means. Differences on day 5 were not significant (p=0.96, Kolmogorov Smirnov test). **e** Tissue titers for colons on day 5 post infection from 2 independent experiments; blue squares represent means + SD. Limit of detection ≈2.5 $\log_{10}CCID_{50}$/g. Differences on day 5 were not significant (p=0.76, Kolmogorov Smirnov test).

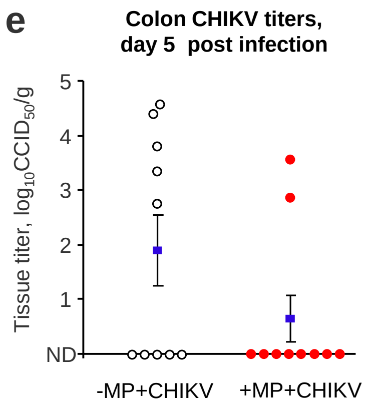

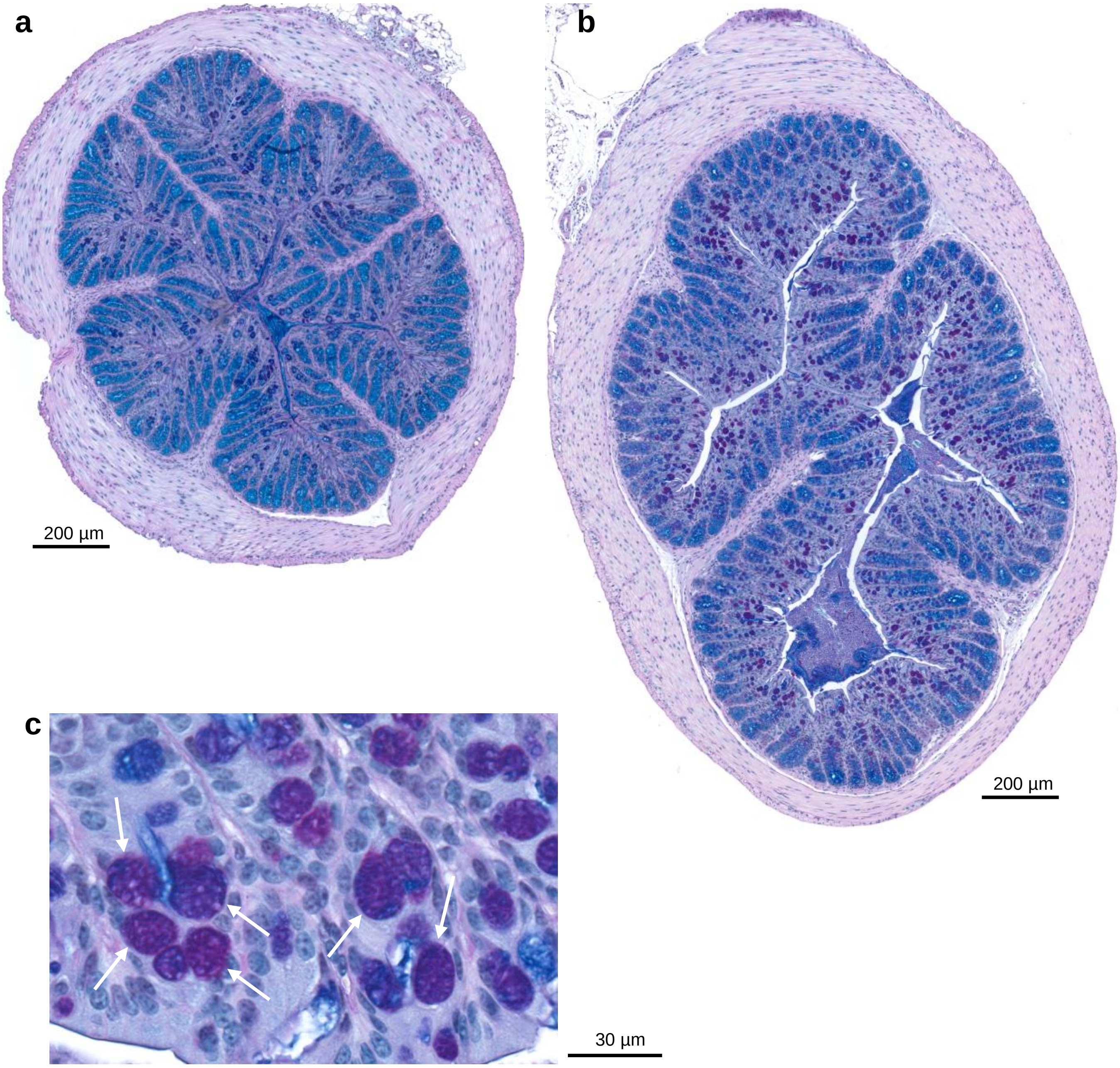


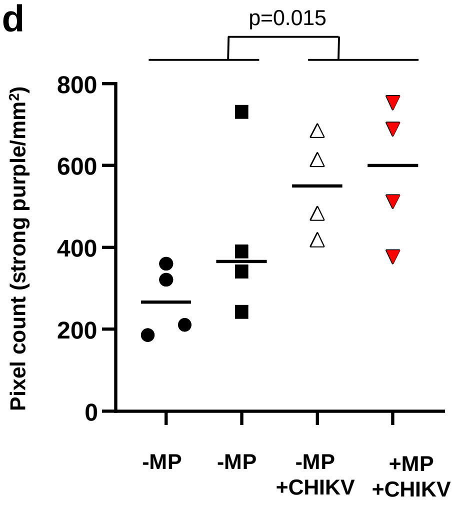


**Supplementary Fig. 5. AB/PAS staining of colon. a-c** Colon from day 9 post infection stained with Alcian blue/periodic acid-Schiff (AB/PAS). **a** +MP. **b** +MP+CHIKV. **c** High resolution image with dark purple staining goblet cells indicated by white arrows. **d**. Aperio pixel count image analysis (n=4 mice per group). Statistics by t test.

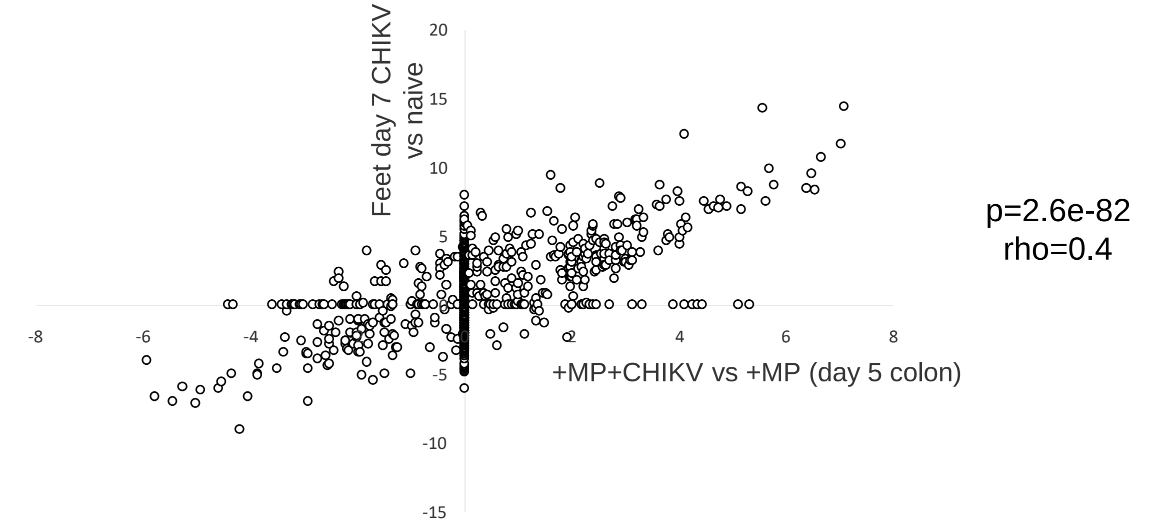


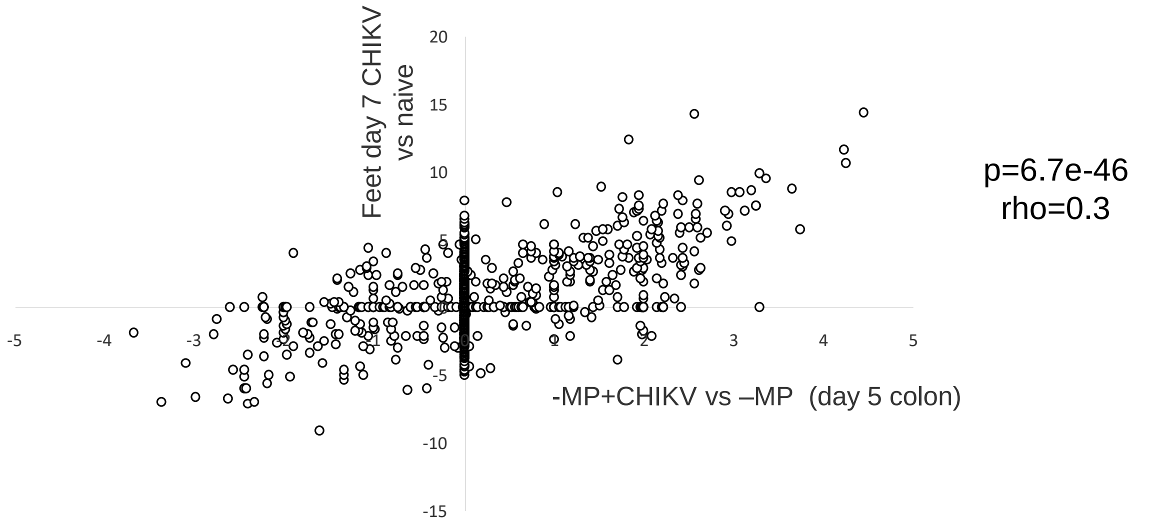


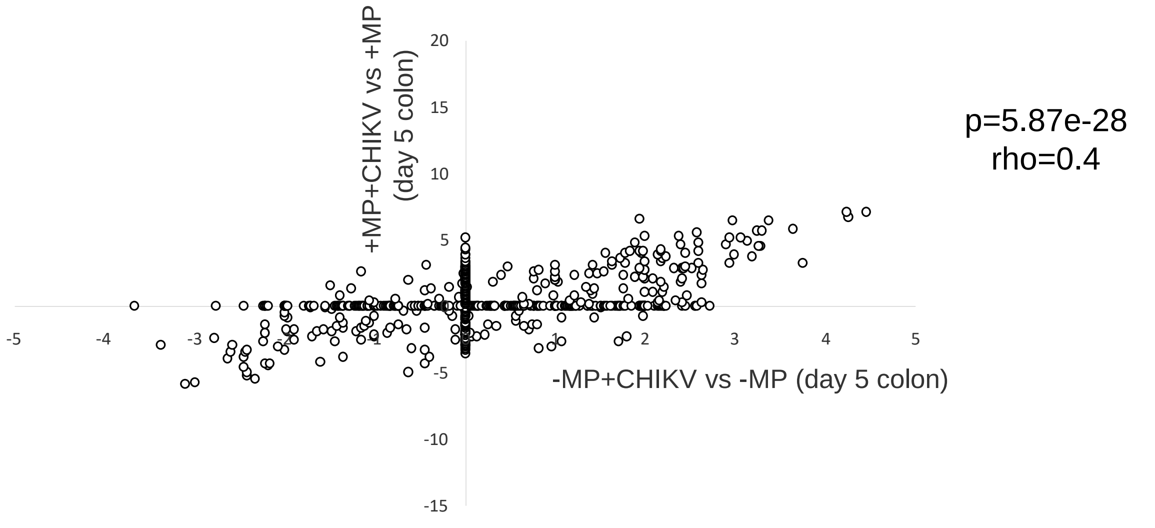


**Supplementary Fig. 6. USR correlations for CHIKV signatures.** The DEGs from Wilson et al 2017 (CHIKV infected day 7 feet vs naïve feet), colon day 5 post CHIKV infection for +MP+CHIKV vs +MP-CHIKV and -MP+CHIKV vs –MP-CHIKV were analysed by IPA upstream regulator (USR) feature (Supplementary Tables 3c, 3f and 3i, respectively). USR Z scores are plotted. Where a USR annotation is provided by analysis of one DEG list, but not the other, the latter is given a nominal z score of 0. Statistics by Spearman correlation.

a

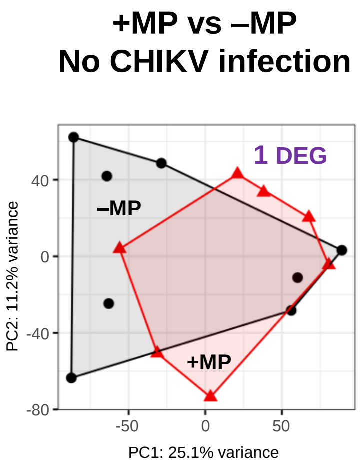


b

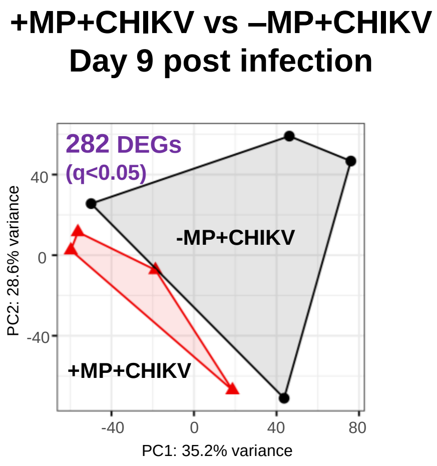


c

| Sample | Total reads | CHIKV reads |
|---|---|---|
| +MP+CHIKV day 9 MLN | 65622286 | 0 |
| +MP+CHIKV day 9 MLN | 54755094 | 0 |
| +MP+CHIKV day 9 MLN | 74696702 | 0 |
| +MP+CHIKV day 9 MLN | 68599238 | 0 |
| -MP+CHIKV day 9 MLN | 70044240 | 0 |
| -MP+CHIKV day 9 MLN | 71584414 | 0 |
| -MP+CHIKV day 9 MLN | 73861786 | 0 |
| -MP+CHIKV day 9 MLN | 62863774 | 0 |

d

| gene | log2FC | gene | log2FC |
|---|---|---|---|
| PPIB | -0.33 | PDZD8 | 0.19 |
| PDIA6 | -0.24 | ACAP2 | 0.19 |
| TLE5 | -0.19 | ITPR1 | 0.19 |
| TPT1 | -0.19 | FOXN2 | 0.20 |
| FAU | -0.18 | SON | 0.20 |
| RPL18 | -0.18 | XIAP | 0.20 |
| RPL5 | -0.17 | CCAR1 | 0.20 |
| MED28 | -0.17 | CFLAR | 0.21 |
| RPL10A | -0.16 | ETV3 | 0.21 |
| RACK1 | -0.15 | ATRX | 0.21 |
| EIF3G | -0.15 | UBR5 | 0.23 |
| PPIA | -0.12 | ASXL2 | 0.23 |
| ITCH | 0.13 | NUP160 | 0.23 |
| FAM91A1 | 0.14 | CBL | 0.24 |
| Zfp148 | 0.16 | AGO3 | 0.24 |
| GOPC | 0.16 | DMXL1 | 0.24 |
| BCLAF1 | 0.16 | PRKX | 0.25 |
| PHF3 | 0.17 | CHD1 | 0.25 |
| CLTC | 0.17 | SYNJ1 | 0.26 |
| NR3C1 | 0.17 | FN1 | 0.29 |
| DHX33 | 0.18 | AGO2 | 0.29 |
| MTR | 0.18 | SPEN | 0.30 |
| RPS6KA3 | 0.18 | EP300 | 0.34 |
| WNK1 | 0.19 | ATM | 0.35 |
| RC3H1 | 0.19 | KAT6A | 0.36 |

e

| GO biological process | Fold Enrichment | FDR (q) |
|---|---|---|
| negative regulation of nitrogen compound metabolic process | 4.74 | 1.08E-05 |
| negative regulation of macromolecule metabolic process | 4.31 | 1.09E-05 |
| negative regulation of cellular metabolic process | 4.44 | 1.67E-05 |
| negative regulation of metabolic process | 4.18 | 1.69E-05 |
| negative regulation of cellular process | 3.01 | 2.62E-05 |
| negative regulation of biosynthetic process | 5.5 | 6.01E-05 |
| regulation of cellular metabolic process | 2.62 | 1.22E-04 |
| negative regulation of biological process | 2.75 | 1.22E-04 |
| regulation of nitrogen compound metabolic process | 2.69 | 1.62E-04 |

**Supplementary Fig. 7. RNA-Seq analyses of MLNs. a** PCA plot for MLNs after 33 and 37 days with (+MP) and without (-MP) MP consumption via the drinking water (mice from two time point were combined). PCA plot of the first two principal components of the log2 normalised counts of all genes that passed independent filtering by DESeq2. The DEG was Hnrnpc. **b** PCA plot for MLNs day 9 post infection. **c** RNA-Seq analysis of MLN day 9 post infection identified no reads from any MLN mouse/sample that mapped to the CHIKV genome. **c** RNA-Seq analysis of MLN day 9 post infection identified no reads from any MLN mouse/sample that mapped to the CHIKV genome.
**d** DEGs from +MP+CHIKV vs –MP+CHIKV day 9 MLN that gave rise to the virus annotations in Supplementary Table 4g (concatenated). **e** UP DEGs in d analysed by Gene Ontology, top annotations by q value shown.

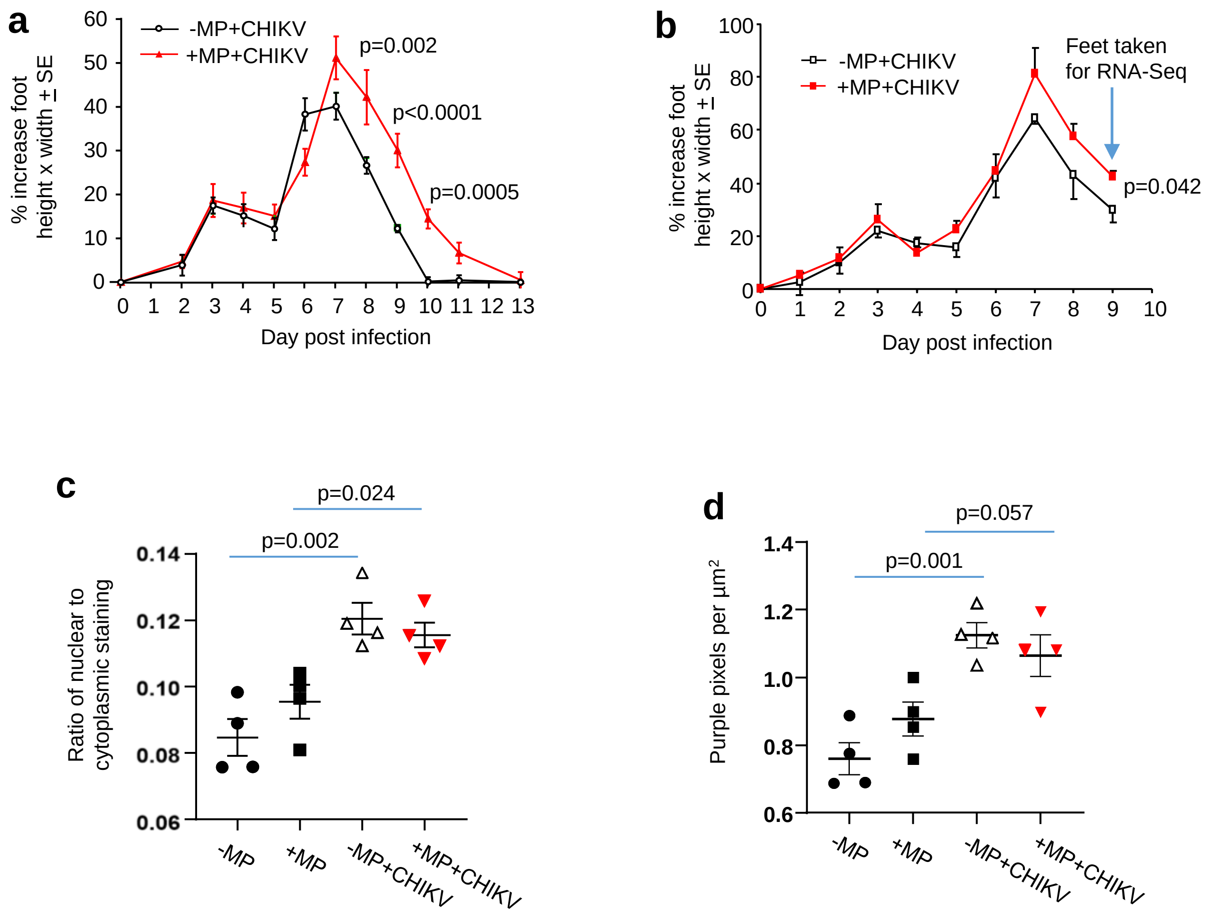


**Supplementary Fig. 8. Foot swelling and cellular infiltrates.** **a** Repeat experiment of that shown in Fig. 4a. n=12 feet from 6 mice per group. Statistics by Kolmogorov-Smirnov tests. **b** Foot swelling of the subset of 4 feet from 4 mice per group harvested for RNA-Seq (Fig. 4a). These mice/feet represent a subset of feet/mice of those feet/mice described in Fig. 4a. Statistics by Kruskal-Wallis test. **c** H&E staining of foot sections from the indicated groups (three replicate sections were stained and scanned using Aperio to provide one mean number for each foot; one foot per mouse, n=4 mice per group). The ratio of blue/purple (nuclei) to red (cytoplasmic) staining represents a simple measure of lymphocyte/leukocyte infiltration, as these cells have a much higher nuclear to cytoplasmic ratio than tissue resident cells. Statistics by t tests. **d** Using the same sections as in c, the nuclear (blue/purple) pixel count per unit area. Statistics by t tests.

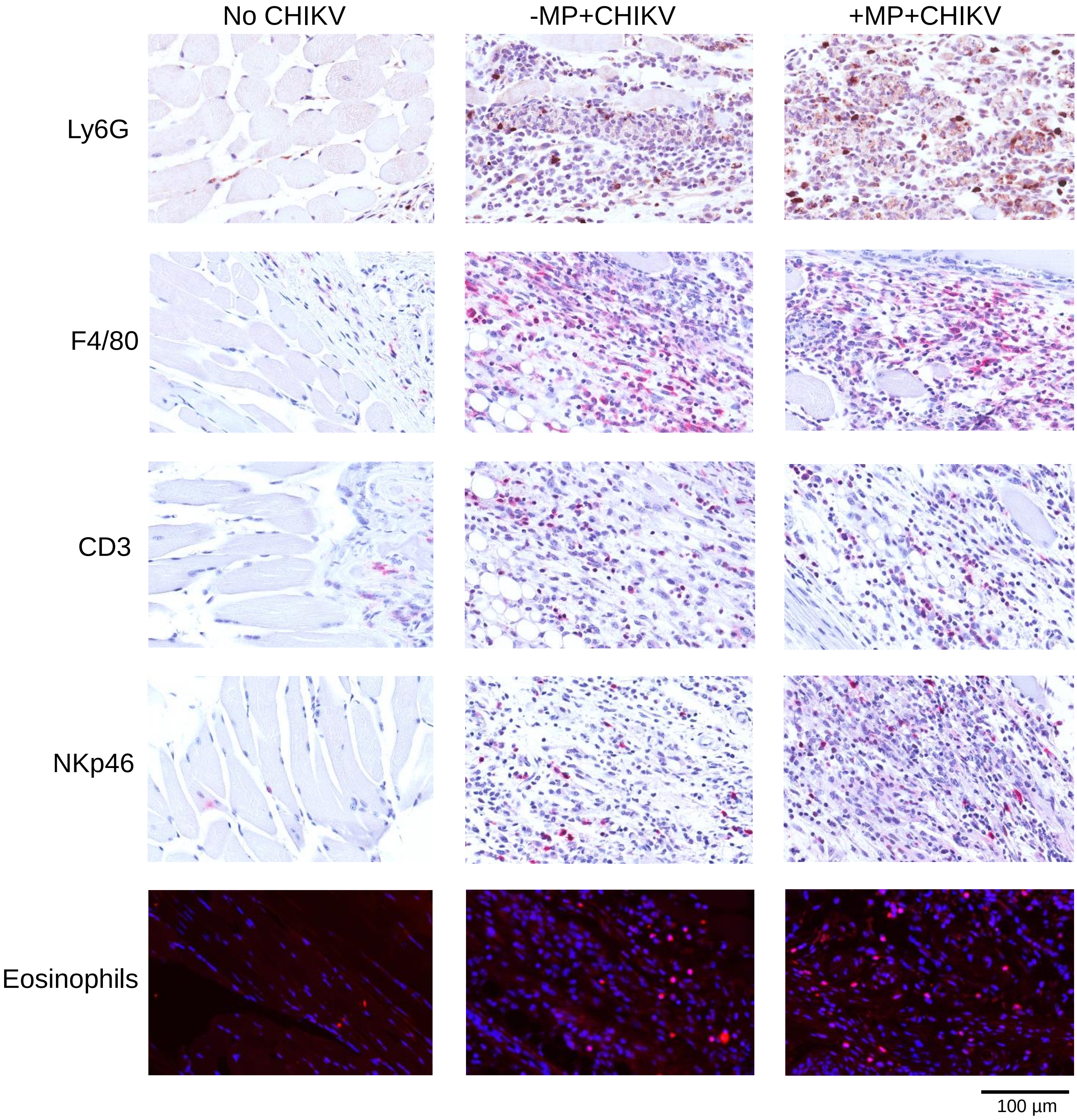


**Supplementary Fig. 9. IHC feet day 9.** Examples of IHC staining of feet day 9 post infection with CHIKV. Ly6G used Nova Red, and F4/80, CD3 and NKp46 used Warp Red, for detection. Akoya Opal 620 tyramide was used for eosinophils.

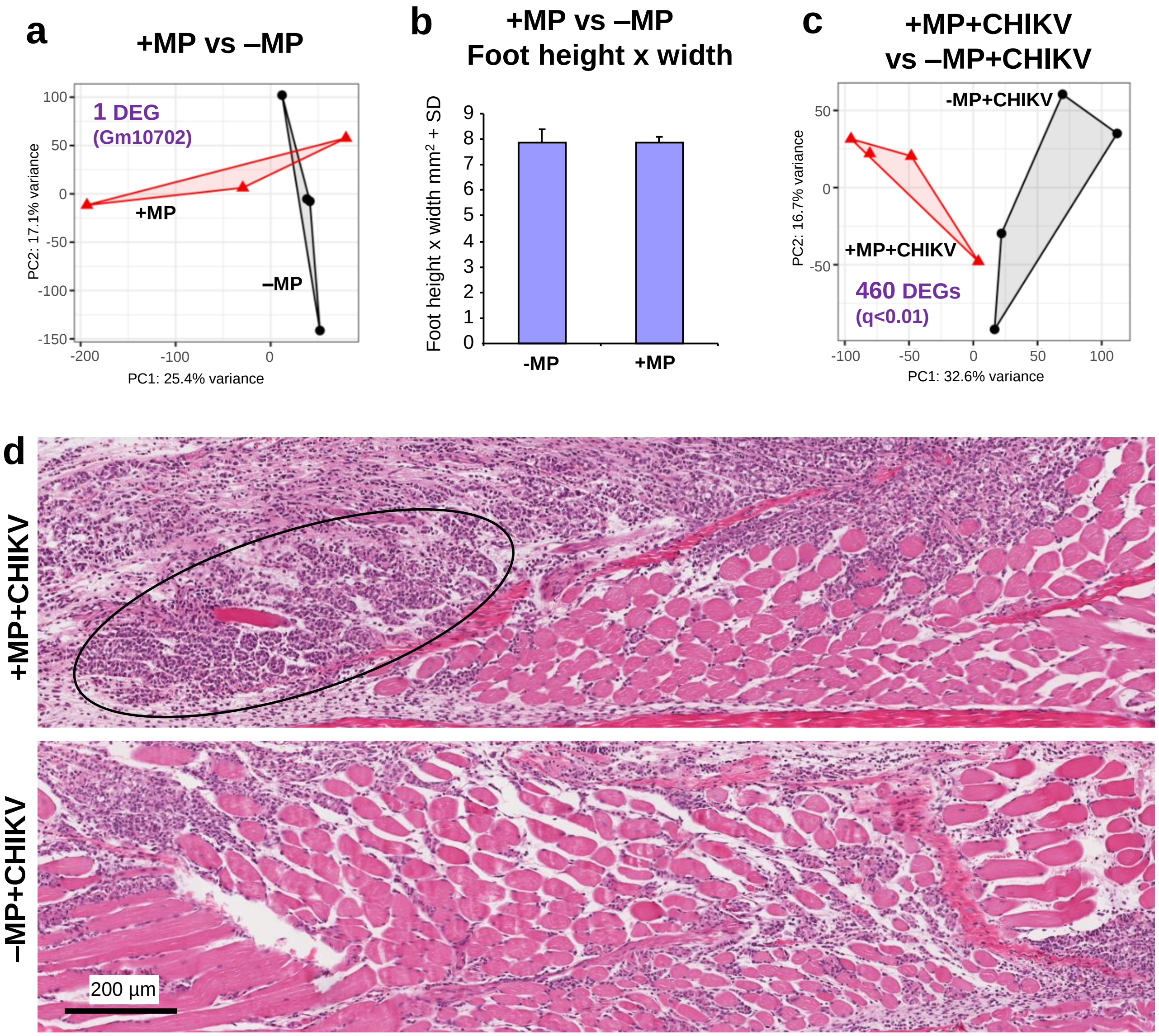


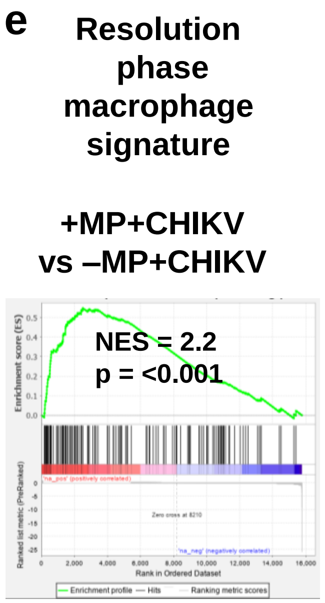


**Supplementary Fig. 10. Feet day 9; +MP vs-MP and +MP+CHIKV vs-MP+CHIKV. a** PCA plot from RNA Seq data using the first two principal components of the log2 normalised counts of all genes that passed independent filtering by DESeq2. Feet taken on day 37 (28 + 9) post onset of MP supplementation; no CHIKV infection.
**b** Foot measurements taken on day 37 (28 + 9) post onset of MP supplementation; no CHIKV infection. n=12 mice and 24 feet per group. **c** PCA plot as in (a) for data from Feet taken on day 37 post onset of MP supplementation and 9 days post infection. **d** H&E staining of muscle in the feet day 9 post infection. Black oval – clear destruction of myocyte bundles and replacement with mononuclear infiltrates.
**e** Genes up-regulated in resolution phase macrophages (Prow et al, 2019) were used in a GSEA against All genes for feet day 9 post infection for +MP+CHIKV vs –MP+CHIKV (Supplementary Table 5a) pre-ranked by fold change. A highly significant enrichment was observed with a positive NES score, arguing that resolution phase macrophages were increased in the +MP+CHIKV feet. As inflammation is increased in this group, an increase in the resolution phase might be expected.

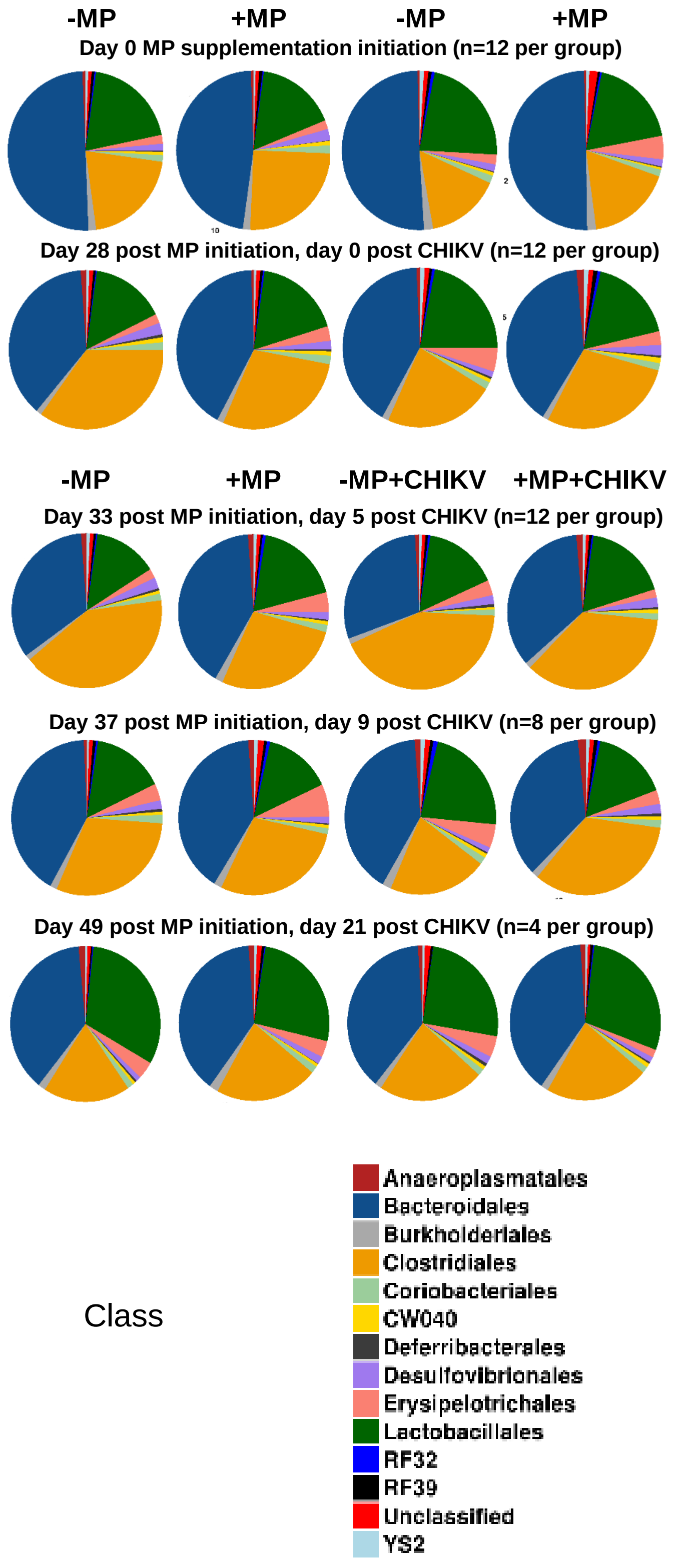


**Supplementary Fig. 11. Microbiome by class**. Arranged as in Fig. 6a**.**

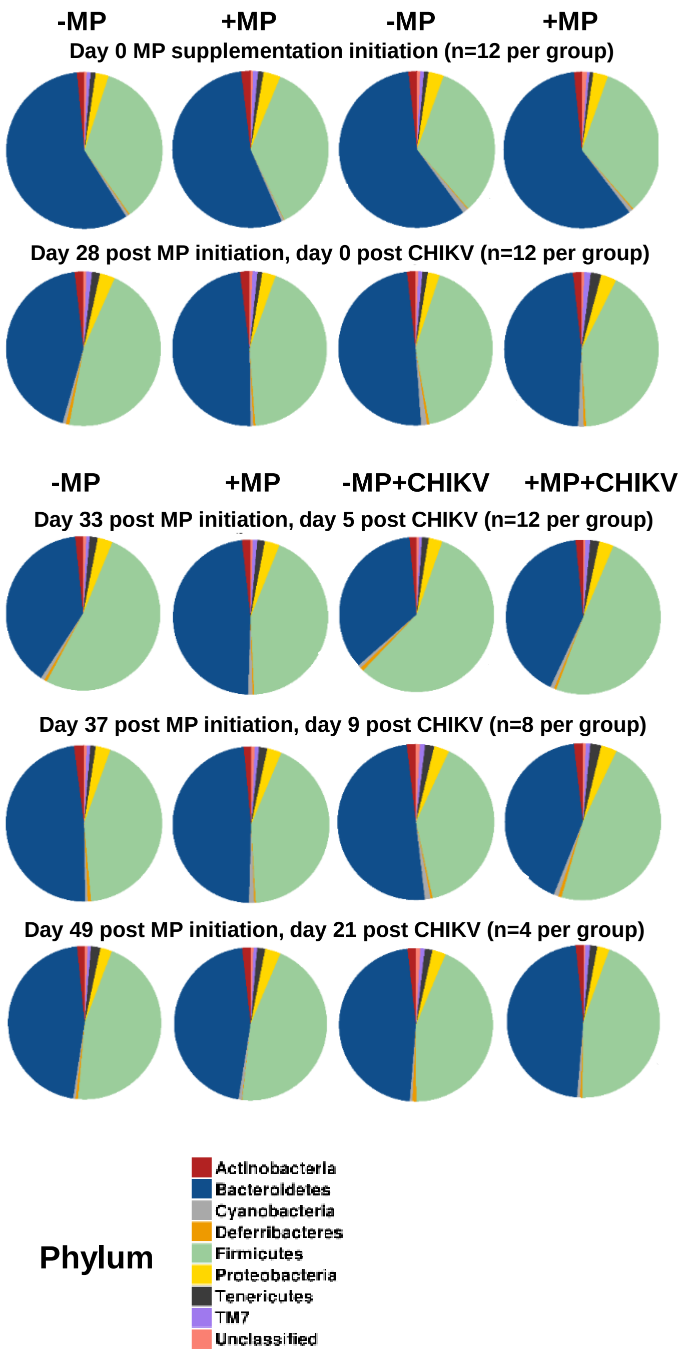


**Supplementary Fig. 12. Microbiome by phylum**. Arranged as in Fig. 6a**.**

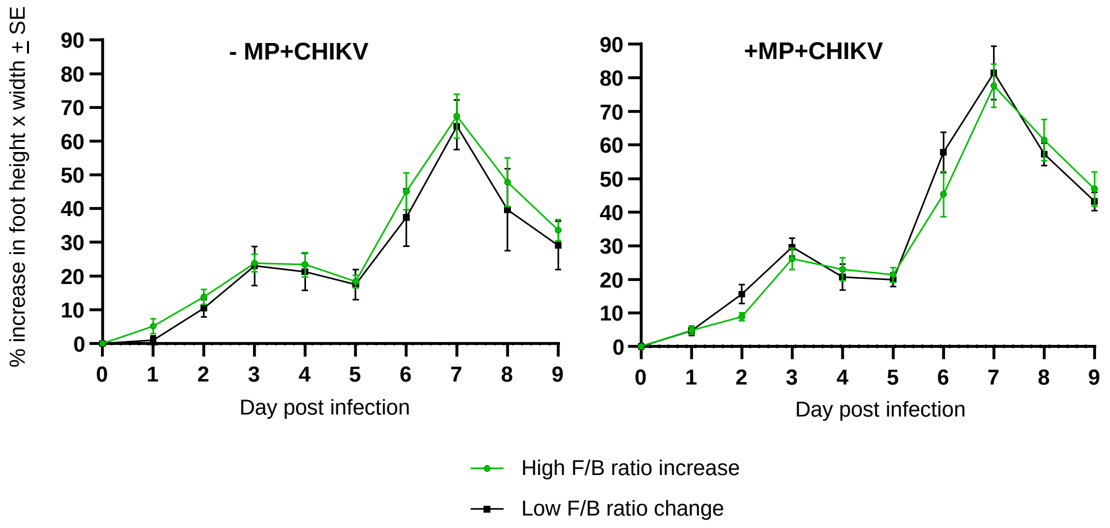


**Supplementary Fig. 13. Foot swelling for mice with increased vs low F/B ratio change mouse groups.** Foot swelling for the two mouse groups described in Fig. 7c.

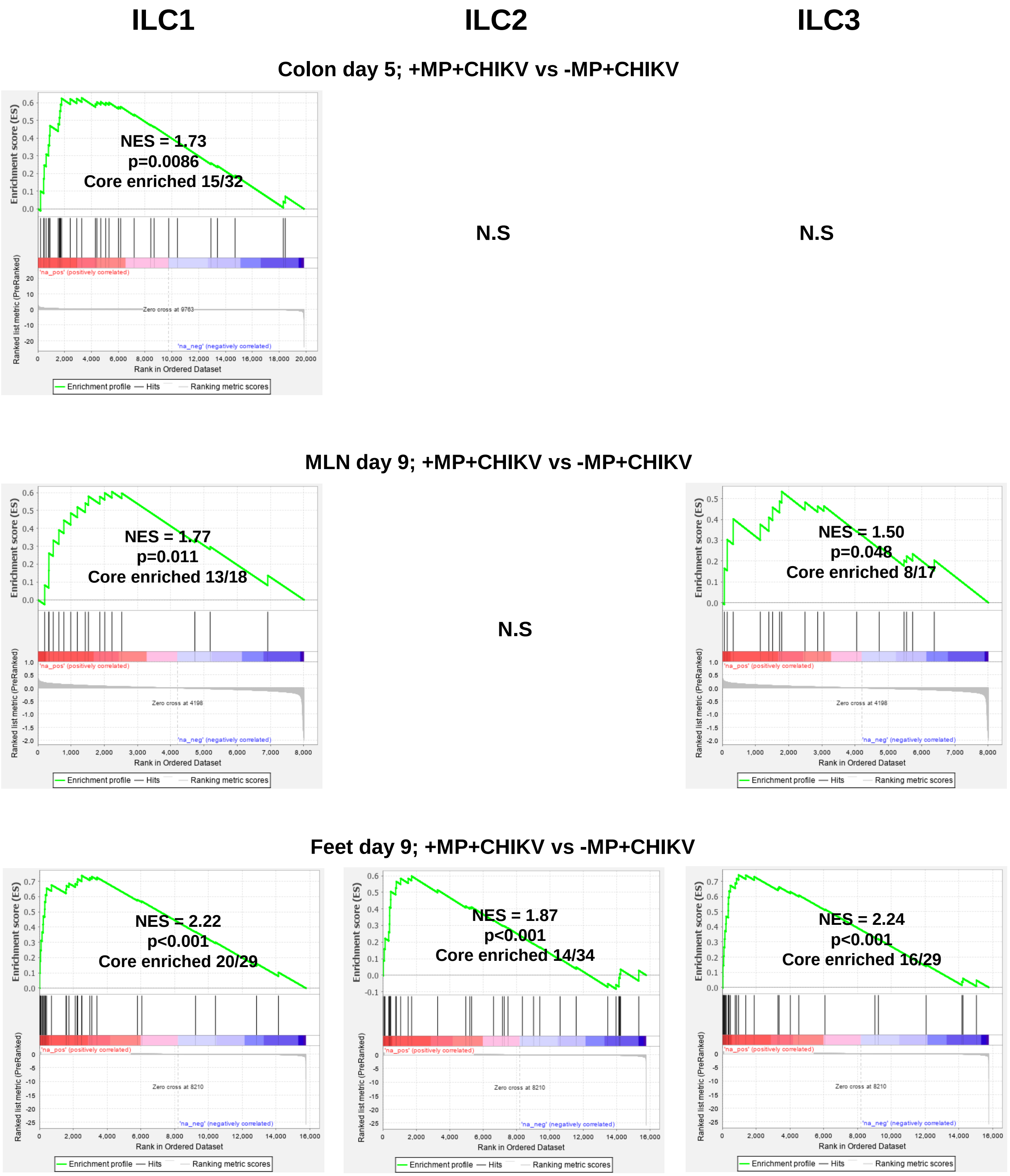


**Supplementary Fig. 14. ILC GSEAs.** GSEAs were undertaken using gene sets constructed from the literature for ILC1, ILC2 and ILC3 (Supplementary Table 7a). These gene sets were used to interrogate all lists from the indicated +MP+CHIKV vs -MP+CHIKV RNA-Seq comparisons pre-ranked by fold change. Core enriched genes are provided in (Supplementary Table 7b). Note the All lists have genes missing were DESeq2 returns an NA for FDR, so the denominator for core enriched genes (e.g. 32, 18 and 29) differs for each GSEA despite use of the the same gene set (e.g. ILC1). N.S. – not significant.

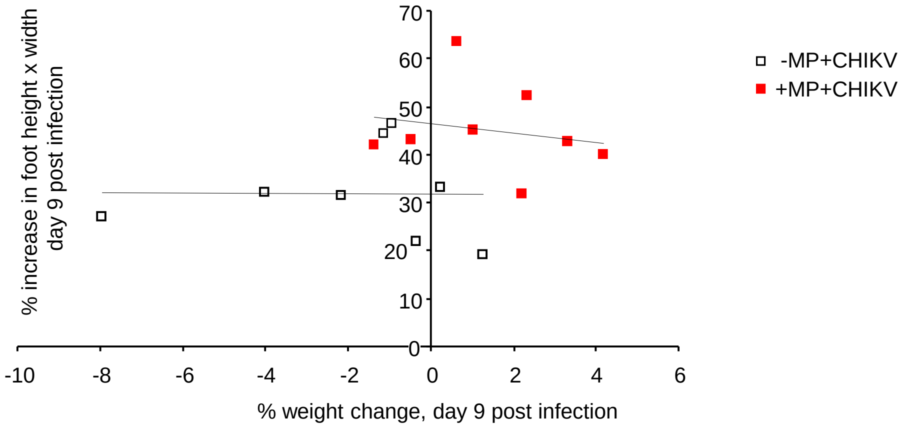


**Supplementary Fig. 15. Weight vs foot swelling day 9.** Individual mice in the +MP+CHIKV and –MP+CHIKV groups plotted for their % foot swelling and weight change on day 9 post infection (both relative to day 0). The two groups are significantly different p=0.0067 (statistics by multivariate ANOVA). Linear trendlines are shown for each group and illustrate that weight change and foot swelling do not correlate, with no indication that increased body weight leads to increased foot swelling.
